# Phylotranscriptomics points to multiple independent origins of multicellularity and cellular differentiation in the volvocine algae

**DOI:** 10.1101/2021.03.16.435725

**Authors:** Charles Ross Lindsey, Frank Rosenzweig, Matthew D Herron

## Abstract

The volvocine algae, which include the single-celled species *Chlamydomonas reinhardtii* and the colonial species *Volvox carteri*, serve as a model in which to study the evolution of multicellularity and cellular differentiation. Studies reconstructing the evolutionary history of this group have often relied on datasets of one to a few genes for phylogenetic inference and ancestral character state reconstruction. These studies suggest that multicellularity evolved only once in the volvocine algae, that each of its three colonial families is monophyletic, and that there have been at least three independent origins of cellular differentiation in the group. We performed RNA-Seq on 55 strains representing 47 volvocine algal species and obtained similar data from curated databases on 13 additional strains. We compiled a dataset consisting of transcripts for 40 single-copy, protein-coding, nuclear genes, then subjected the predicted amino acid sequences of these genes to maximum likelihood, Bayesian inference, and coalescent-based analyses. These analyses show that multicellularity independently evolved at least twice in the volvocine algae and that the colonial family Goniaceae is not monophyletic. Our data further indicate that cellular differentiation independently arose at least four and possibly as many as six times within the group. Altogether, these results show how multicellularity and cellular differentiation are evolutionarily labile in the volvocine algae, affirming their importance for the study of major transitions in the history of life.

## Introduction

The evolution of multicellularity is widely considered a major transition in the history of life (Grosberg & Strathmann, 2007; Maynard Smith & Szathmáry, 1995; Szathmáry & Maynard Smith, 1995; West *et al*., 2015). Multicellularity not only gave rise to most of the visible life forms on the planet, but also opened the door to cellular differentiation, including that between somatic and reproductive cells, a hallmark feature of sexual reproduction in eukaryotes that exhibit morphological complexity (Hallmann, 2011; Szathmáry & Maynard Smith, 1995; Wahl & Murray, 2016). Questions regarding the evolution of multicellularity and cellular differentiation have been approached using the fossil record (Chen *et al*., 2014; Han & Runnegar, 1992; Schirrmeister *et al*., 2013), laboratory evolution (Herron *et al*., 2019; Quintero-Galvis *et al*., 2018; Ratcliff *et al*., 2012, 2013), and comparative approaches that include superimposing cell biology upon molecular phylogeny (Herron & Michod, 2008; Kirk, 2005; Koyanagi, 2015). The last of these approaches is predicated on the assumption that the cell biology and molecular phylogeny are mutually informative, an assumption that requires the phylogeny itself to be accurate.

The volvocine green algae have proved especially useful for investigating the major transition leading to multicellularity. The group consists of ∼50 extant species which exhibit a range of body plans, cell numbers, sizes, and forms of sexual reproduction. The smallest of these are single-celled (e.g., *Chlamydomonas reinhardtii*); the largest, at up to 3 mm in diameter and up to 50,000 cells, are spheroidal, swimming colonies in the genus *Volvox*. Since the initial “very pleasant sight” of swimming *Volvox* colonies described by Van Leeuwenhoek more than 300 years ago (van Leeuwenhoek, 1700), the volvocine algae have come to be accepted as a useful model system in which to address questions related to the origins of multicellularity and cellular differentiation (Kirk, 2001; Schmitt, 2003). Multiple species have now had their genomes sequenced (Craig *et al*., 2021; Featherston *et al*., 2018; Hanschen *et al*., 2016; Merchant *et al*., 2007; Prochnik *et al*., 2010), and those of unicellular *C. reinhardtii* and multicellular *V. carteri* forma *nagariensis* are well-annotated (Merchant *et al*., 2007; Prochnik *et al*., 2010). However, the volvocine algae encompass more than two organisms representing alternative forms of life in terms of size and development. Vegetative forms range in characteristic cell number from 1 to ∼50,000 and exhibit intermediate degrees of complexity likely similar to extinct ancestors. Further, multicellularity and cellular differentiation arose within the volvocine algae much more recently than those traits arose in animals: ∼220 million years ago (Herron *et al*., 2009) versus ∼600 million years ago (Brunet & King, 2017), respectively.

Evolution of the volvocine algae has sometimes been viewed as a linear progression in size and complexity (Lang, 1963; Larson *et al*., 1992). Unicellular taxa such as *Chlamydomonas* occupy one end of this continuum, while fully differentiated, multicellular taxa such as *Volvox* occupy the other. This concept, the ‘volvocine lineage hypothesis’, used a streamlined phylogeny of the volvocine algae to help explain how a multicellular species with complete germ-soma differentiation such as *Volvox* might evolve from a unicellular, *Chlamydomonas*-like ancestor. However, morphological and molecular phylogenetic studies suggest that the history of the volvocine algae may be more complicated, as cellular differentiation, different modes of sexual reproduction, and varying body plans appear to have evolved multiple times within the group (Hanschen *et al*., 2018b; Herron *et al*., 2010).

Current understanding of the major evolutionary relationships within this group has often been based on the analysis of five chloroplast gene sequences (Hanschen *et al*., 2018; Herron *et al*., 2009; Herron & Michod, 2008; Nakada *et al*., 2010; Nozaki *et al*., 2000, 2014, 2015). Chloroplast gene-based phylogenies have also been used to carry out ancestral-state reconstructions (Grochau-Wright *et al*., 2017; Hanschen *et al*., 2018; Herron *et al*., 2010; Herron & Michod, 2008), opening a window on how multicellularity and cellular differentiation evolved within the volvocine algae. Overall, the branching order of most chloroplast gene-based phylogenies is defined by two related groups: *(i)* a set of unicellular species (e.g., *Chlamydomonas reinhardtii*) that are paraphyletic with respect to *(ii)* a clade that encompasses the three major families of colonial volvocine algae: Tetrabaenaceae (*Tetrabaena* and *Basichlamys*), Goniaceae (*Gonium* and *Astrephomene*), and Volvocaceae (*Colemanosphaera, Eudorina, Pandorina, Platydorina, Pleodorina, Volvox, Volvulina,* and *Yamagishiella*) (**Figure 1A** and **1D**). In this scheme, the Tetrabaenaceae is a sister group to the clade formed by the Goniaceae and Volvocaceae. Although this framework only takes into account family-level relationships, several conclusions can be drawn. *First*, the colonial species form a clade. *Second,* each of the three families is monophyletic. *Third,* monophyly among the colonial species implies that multicellularity evolved only once within that group with no reversion to unicellularity.

**Figure 1.**
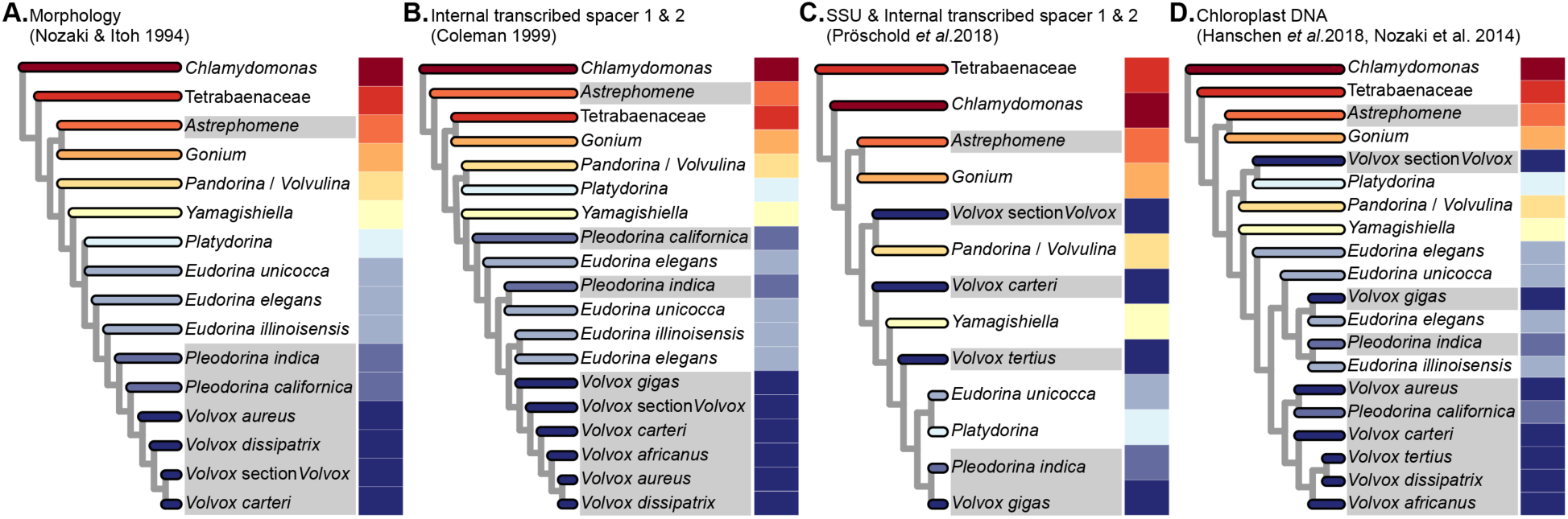
**Phylogenies of the volvocine algae are not concordant:** Four volvocine green algae phylogenies based on different types of data, displayed in chronological order of their appearance in the literature. Species highlighted in shades of gray exhibit somatic cell differentiation. The varying colors to the right of each phylogeny have been arbitrarily assigned to particular genera and are intended to be used as a visual aid to highlight differences among the phylogenies.

Two recent studies have called into question the monophyly of the colonial volvocine algae (**Figure 1C**). Pröschold *et al*. (2018) based their inferences on two datasets: one consisting of SSU *r*DNA sequences plus internal transcribed spacer (ITS) sequences 1 and 2, the other consisting of ITS sequences alone. Nakada *et al*. (2019) used a single-gene 18S *r*RNA dataset. Both studies inferred that the colonial species are paraphyletic with respect to certain unicells in the genera *Chlamydomonas* and *Vitreochlamys*.

The taxonomic status of the Goniaceae has also been called into question by studies (**Figure 1B**) that indicate the group is either not monophyletic (Coleman, 1999; Nozaki *et al*., 2000) or that there is low support for a sister relationship between *Astrephomene* and *Gonium* (Nakada *et al*., 2019; Nozaki *et al*., 2000; Pröschold *et al*., 2018). Moreover, a number of recent volvocine algal phylogenies leave uncertainty as to how many times cellular differentiation evolved within the group. Chloroplast sequence data suggest at least 3 independent origins of cellular differentiation: in *Astrephomene*, in *Volvox* section *Volvox* (sometimes referred to as *Euvolvox*), and in the *<u>E</u>udorina, <u>V</u>olvox, <u>P</u>leodorina* (EVP) clade (**Figure 1B-D**). Within the EVP clade it is unclear whether cellular differentiation in *Pleodorina thompsonii, Volvox gigas* and *V. powersii,* and *Pleodorina starrii* and *P. indica* arose independently from that in *V. carteri* (**Figure 1B-D**).

The foregoing uncertainties highlight the need for a new and more robust molecular phylogeny of the volvocine algae. These uncertainties may arise from incomplete taxonomic sampling, limited genetic sampling, or both. While five volvocine algal species have had their genomes sequenced, most taxonomically comprehensive phylogenetic inferences about this evolutionarily important group have been constructed using relatively small datasets. These consist of either the sequence of five chloroplast genes (Hanschen *et al*., 2018; Herron & Michod, 2008; Hu *et al*., 2019; Nozaki *et al*., 2014) representing an aggregate of ∼6000 nucleotide positions. Others consist of small (≤6) multi-gene datasets consisting of chloroplast gene(s), ribosomal molecular markers, or both (Pröschold *et al*., 2018, Nakada *et al*., 2019). Moreover, the use of chloroplast genes in phylogenetic reconstruction can be problematic because they are effectively a single linkage group, they vary little among recently diverged species (Dong et al., 2012), and they are at increased risk of incomplete lineage sorting due to the retention of ancestral polymorphisms (Jakob & Blattner, 2006; Xu *et al*., 2012).

Of special concern is the observation that volvocine phylogenies inferred using chloroplast genes **(Figure 1D)** conflict with those constructed using nuclear genes (**Figures 1B & 1C**) (Coleman, 1999; Nakada *et al*., 2019; Pröschold *et al*., 2018). While conflicts between chloroplast and nuclear phylogenies are not unusual (Rose *et al*., 2020; Soltis & Kuzoff, 1995; Yu *et al*., 2013), they do foster ambiguity.

Here, we seek to resolve volvocine relationships using taxonomically dense sampling of multiple, unlinked loci. We have adopted a phylotranscriptomic approach that uses a concatenated amino acid alignment of 40 nuclear protein-coding, single-copy genes. We sequenced whole transcriptomes of 55 strains encompassing 47 nominal species and used previously published RNA-Seq data for 9 strains and shared amino acid sequences for 4 strains. Our goal was to derive a robust phylogeny of the volvocine algae that would enable inferences about the evolution of multicellularity, cellular differentiation, sexual dimorphism, and other traits in this group. Our results represent the most taxonomically comprehensive phylogeny yet produced of the volvocine algae using a nuclear dataset, including all described genera and multiple representatives of all genera that are not monotypic. Our results show that the colonial species do not form a clade, that the Goniaceae are not monophyletic, and that multicellularity has independently evolved at least twice and cellular differentiation at least four times within the volvocine algae.

## Materials and Methods

### Strains and culture conditions

Algal strains used in this study were obtained from the National Institute for Environmental Studies (NIES, Japan), the Culture Collection of Algae at the University of Göttingen (SAG, Germany), and the Culture Collection of Algae at the University of Texas at Austin (UTEX, USA). Strain provenance and culture collection ID numbers are shown in **Table 1**, with previously published data designated with an asterisk. All cultures were grown at 20-26^°^C under cool-white LED lamps (4300K) with an intensity of 2500-2700 lux under a 14-hour light/10-hour dark cycle. A detailed description of the medium used to culture each strain, as well each strain’s morphology, degree of cellular differentiation, and gamete size is provided in **Supplementary Materials: Tables S1 and S2, respectively.**

**Table 1.**
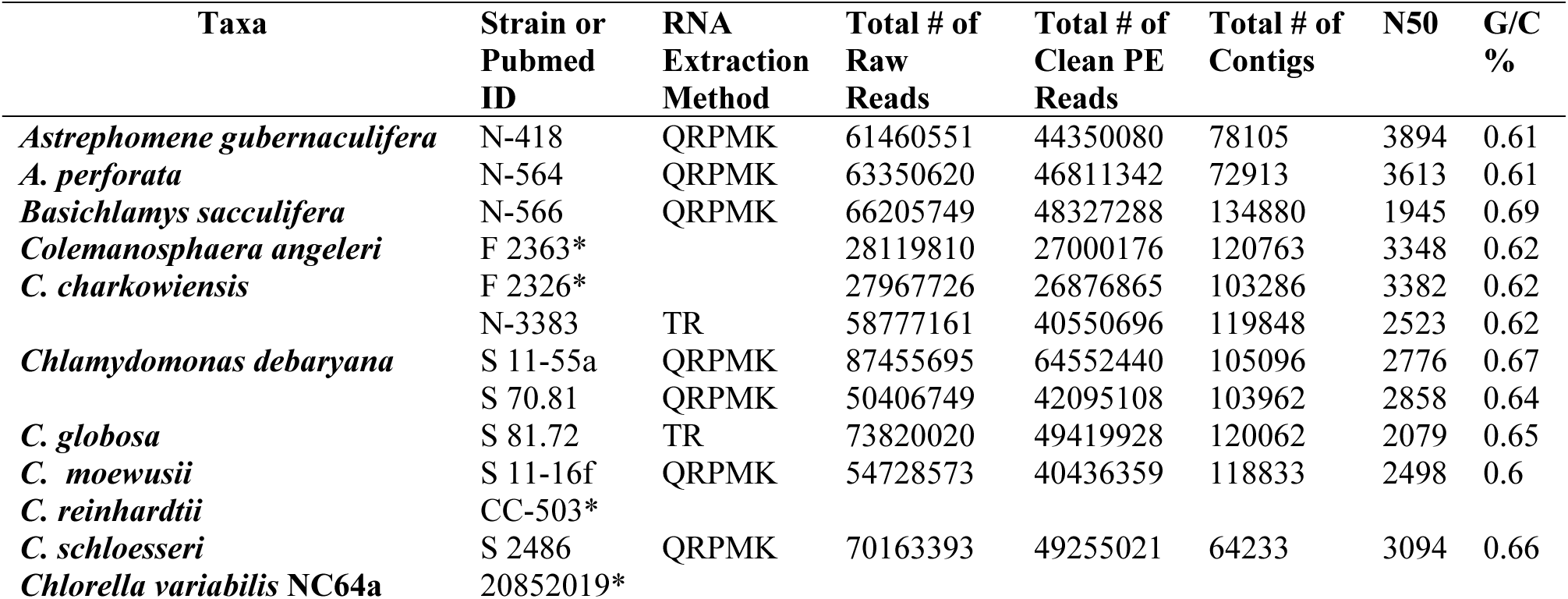
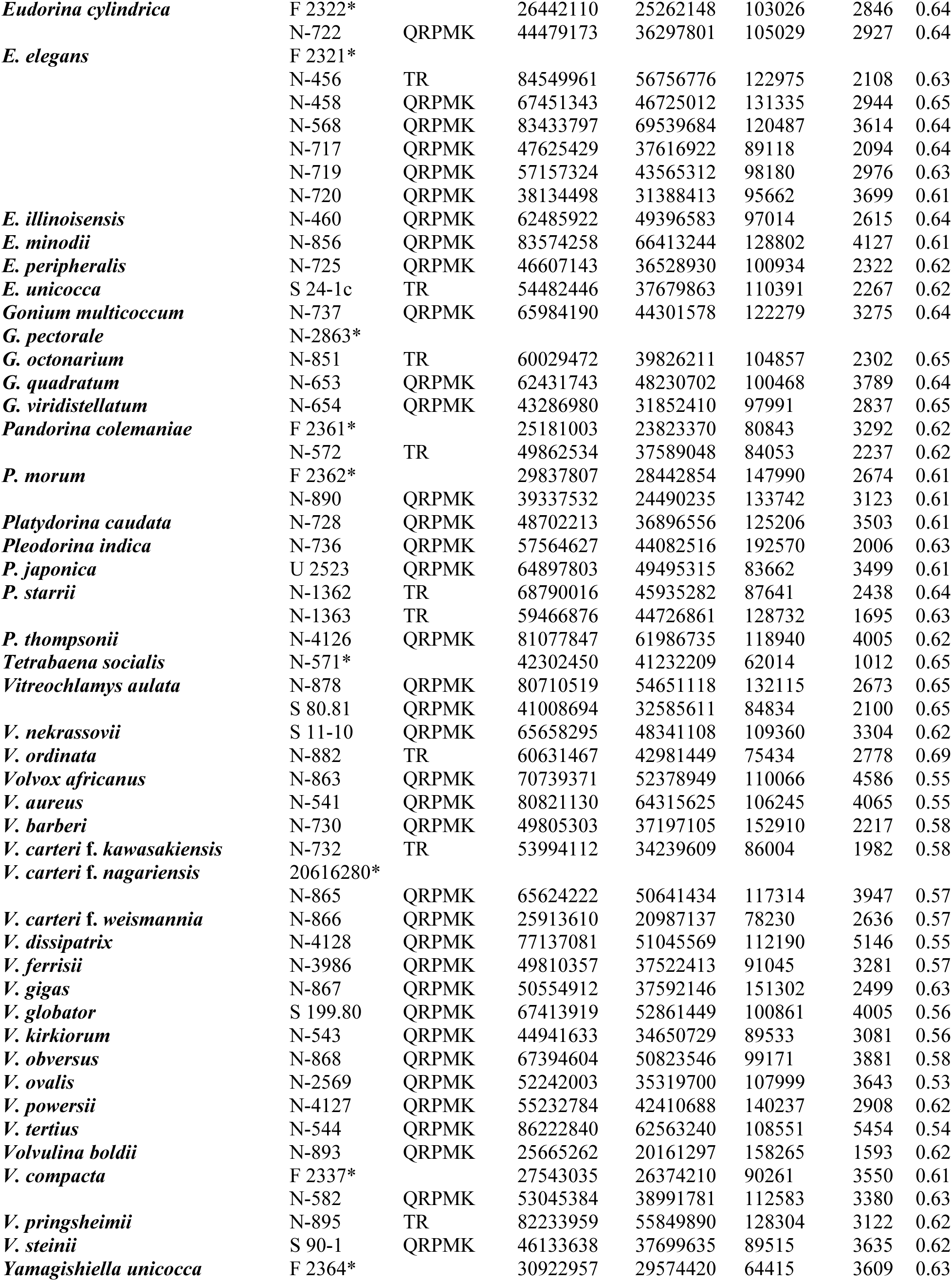
List of taxa used in this study and summary of sequencing and assembly. Under Strain or Pubmed ID, “CC” refers to Chlamydomonas Culture Collection at the University of Minnesota (CC, USA), “F” refers to Culture Collection of Freshwater Algae at the Institute of Hydrobiology, Chinese Academy of Sciences (FACHB, China), “N” refers to National Institute for Environmental Studies (NIES, Japan), “S” refers to Culture Collection of Algae at the University of Göttingen (SAG, Germany), and “U” refers to The Culture Collection of Algae at the University of Texas at Austin (UTEX, USA). “QRPMK” and “TR” under RNA Extraction Method refer to QIAGEN RNeasy Plant Mini Kit and TRizol RNeasy, respectively. Strains assigned an asterisk represent data from previously published studies, with accession numbers shown in **Supplementary Materials: Table S3**.

### RNA extraction procedures

Two protocols were used to isolate total RNA: a modified version of the TRizol RNeasy method described by Matt & Umen (2018) and a slightly modified QIAGEN RNeasy Plant Mini Kit protocol. For a detailed description of each please see **Supplementary Materials: File lindseyetal_rna_extr_protocols.docx**. Information on the protocol was used for each strain is provided in **Table 1.**

### Library Preparation and Sequencing

Before generating a sequencing library, RNA quality and quantity were assessed by Nanodrop and Qubit (Thermo Fisher Scientific, Waltham, MA 02451 USA). RNA integrity was evaluated using an Agilent 2100 Bioanalyzer (Agilent Technologies, Santa Clara, CA 95051 USA. mRNA was isolated using poly T beads, whereafter Illumina libraries were prepared using the NEBNext Ultra II Directional RNA Library Prep Kit. Library concentrations were determined fluorometrically; sequencing was carried out on the Illumina NovaSeq 6000 platform (Illumina, Inc., San Diego, CA 92122 USA) to generate 151 bp paired-end reads.

### Quality Control of Reads

Raw read quality was assessed through FastQC v.0.11.8 with an additional FastQC assessment post-trimming. Quality control of the raw reads was completed with Trimmomatic v.0.39 (Bolger *et al*., 2014) where the bases at the 5’ and 3’ end of each read are trimmed if found to be below a quality score of 3. A 4-base sliding window approach was used to trim the rest of the read once average quality fell below a score of 15; reads that were below a minimum length of 36 bases were discarded (LEADING:3 TRAILING:3 SLIDINGWINDOW:4:15 MINLEN:36). If adapter content was detected by FastQC the additional ILLUMINACLIP step was used with the ‘TruSeq3-PE-2.fa’ file provided by the Trimmomatic developers. If performed, the following ILLUMINACLIP parameters were used: 2:30:10 at the beginning of each command line. This allows for 2 “seed” mismatches where the seed is a short segment of the adapter that is being aligned in every section of the read. If more than 2 mismatches occurred, no trimming of the read occurred. Additionally, there had to be at least 30 matched bases in the paired-end palindrome read alignment and at least 10 matched bases between an adapter sequence and read.

### De Novo Assembly

SOAPdenovo-Trans v1.0.4 (Xie *et al*., 2014) was used to assemble *de novo* transcriptomes from the quality filtered, paired-end reads using a k-mer size of 25 (SOAPdenovo-Trans-31mer all -s <config input file> -o <outfile> -K 25). GapCloser from the SOAPdenovo package was utilized to close gaps in each transcriptome using the same configuration file, which contains read-specific information and file paths, from the previous step (-b <config file> -a <.scafSeq file output by SOAPdenovo-Trans> -o <outfile> -l <max read length, int value> -t <thread number>). Default parameters were used for CD-HIT v4.8.1 (Fu *et al*., 2012) to reduce redundant transcripts from our *de novo* transcriptomes.

### Orthologous Gene Identification for Phylotranscriptomic Analysis

The evolutionary history of the volvocine algae dates back at least 200 million years (Herron *et al*., 2009). Over this timescale nucleotide sequences become saturated with substitutions, diminishing their phylogenetic utility (Hasegawa & Hashimoto, 1993). Amino acid sequences were therefore chosen for our alignments, as they are known to be more reliable for ascertaining distant evolutionary relationships (Loomis & Smith, 1990). De Clerck and colleagues identified 58 nuclear protein-coding, single copy genes that were members of highly conserved gene families across the green algae (*Chlorophyceae, Prasinophytes,* and *Trebouxiophyceae*) and land plants (*Streptophyta*) (De Clerck *et al*., 2018). Their amino acid alignment of the 58 nuclear protein-coding genes that includes *Chlamydomonas reinhardtii* CC-503*, Chlorella variabilis* NC64A*, Gonium pectorale* NIES-2863*, and Volvox carteri* HK10 was kindly shared with our research team. Out of the 58 genes shared, we used 40 for our gene alignments. In order to identify those specific genes in the *de novo* transcriptomes of our taxa a Basic Local Alignment Search Tool (BLAST) server was established in our lab, and a unique BLAST database for each taxon was created following the instructions in the BLAST manual. A BLASTP search using the *C. reinhardtii* CC-503*, G. pectorale* NIES-2863, and *V. carteri* HK10 genes from De Clerck *et al*. as our query sequences enabled us to identify the orthologous genes for each of our taxa.

### Gene Sequence Alignments and Phylotranscriptomic Analysis

The BLASTP results were used to identify the scaffold and open read frame where each gene was located in a strain’s transcriptome. Using a proprietary Python script (**Supporting Materials**: **File translate_scaffold.py)** each scaffold was extracted from its transcriptome and translated in the appropriate reading frame, then the translated scaffold was added to an alignment file. For consistency, we generated *de novo* transcriptomes since we lacked a reference genome for most of our sequenced strains. At times, a gene was found to be incomplete for a given taxon due to assembler or sequencing error after manual examination. When this was determined to be the case, the gene was manually stitched together. This was done in a highly conservative manner: if we could not ascertain whether or not a gene was incomplete due to assembler or sequencing error then it was excluded from the alignment for the given species. We treated the data from previously published studies in the same fashion as data generated in our lab by filtering the raw reads through quality trimming, then assembling *de novo* transcriptomes using the same programs and parameters (**Table 1**).

Amino acid sequences were aligned using MUSCLE v3.8.31 (Edgar, 2004). Alignments were also subjected to manual alignment in Aliview v1.26 (Larsson, 2014); extraneous data were trimmed, leaving only the aligned genes. Ambiguously aligned regions were eliminated from each alignment leaving only conserved and reliably aligned regions for phylogenetic analysis using the following parameters in Gblocks v0.91b (Castresana, 2000): -t=p -b3=8 -b4=2 -b5=h - b6=y. Phyutility v2.7.1 (S. A. Smith & Dunn, 2008) was used to concatenate all gene alignment files.

Single-gene alignments were subjected to ML and BI analyses in order to infer single-gene phylogenies. Single-gene phylogenies were then further analyzed using a coalescent-based approach. The concatenated multi-gene alignment was partitioned so that the appropriate model of protein substitution was applied to each gene for the supermatrix phylogenetic approach under ML and BI.

The ML and BI analyses of the concatenated dataset used a partitioning strategy where the best evolutionary model for each gene was predicted by ProtTest v3.4.2 under the Akaike Information Criterion (AIC). For information regarding each predicted evolutionary model please refer to **Supplementary Materials: Table S4**. The ML analysis was conducted using IQtree v1.6.12 (Nguyen *et al*., 2015) under partition models (Chernomor *et al*., 2016). Support values reported for the IQtree ML analysis were estimated through the bootstrap technique where 1,000 ultrafast bootstrap replicates were generated (Hoang *et al*., 2018). The BI analysis was performed with MrBayes 3.2.7a (Ronquist *et al*., 2012) with 3 heated and 1 cold Markov chains, where trees were sampled every 1,000 generations for a total of 1,000,000 generations with 1,000 trees discarded at the beginning of each chain (ngen =100000000, samplefreq=1000, burnin=1000, nruns=4, nchains=4, starttree=random).

ASTRAL (C. Zhang *et al*., 2018) was used to perform the coalescent-based analysis where all 40 single-gene phylogenies (**Supplementary Materials: File lindseyetal_single_gene_phylogenies**) produced by RAxML v8.2.12 (Stamatakis, 2014) were used as the input after collapsing branches with low bootstrap support (<10) using Newick Utilites v1.6 (Junier & Zdobnov, 2010). Posterior probabilities were assessed for the Bayesian and coalescent-based analyses in MrBayes and ASTRAL, respectively. Lastly, Approximately Unbiased (AU) tests with 100,000 RELL re-samplings were conducted to test certain key topologies and hypotheses using IQtree (-zw 100000 -au) (**Supplementary Materials: Figure S1**).

## Results and Discussion

### De novo transcriptome data makes possible 40 single-gene alignments

We sampled 68 taxa representing all presumed major lineages of the colonial volvocine algae and 9 of their nearest unicellular relatives. Because the phylogenetic position of *Chlamydomonas reinhardtii* has recently been called into question (Nakada et al., 2019; Pröschold et al., 2018), we used a member of the Trebouxiophyceae, *Chlorella variabilis*, as an outgroup (**Table 1**). All described volvocine genera were included, with multiple species represented for every genus that is not monotypic. Truly comprehensive taxon sampling was not possible, since several described species, especially in the genus *Volvox*, are no longer available in culture collections. While our main focus was to resolve relationships within the colonial volvocine algae, our study included several closely related unicellular taxa from the genera *Chlamydomonas* and *Vitreochlamys* in order to provide better phylogenetic resolution of the volvocine algae.

The total number of raw reads generated from RNA-sequencing for each species ranged from 25,665,262 to 87,455,695 reads with an average of 60,194,849 reads per species. After quality trimming of the raw reads (see Methods), the total number of clean paired-end reads ranged from 20,161,297 to 69,539,684 with an average of 44,416,935 reads per species (**Table 1**). From the RNA-seq data, we assembled a total of 40 single-gene alignments that were later concatenated to a single alignment representing an aggregate of 12,650 amino acids, equivalent to 37,950 nucleotide positions, with a total of 5992 parsimony-informative sites. Numbers of informative positions in the single-gene alignments ranged from 40 to 446. Trees inferred using maximum likelihood (ML), Bayesian inference (BI), and coalescence-based (CB) analyses were generally well-supported with some topological differences between the ML and BI analyses relative to the CB analysis, as described below.

### Our results conflict with prior volvocine algal phylogenies in four respects

*First*, we find that the colonial volvocine algae are paraphyletic with respect to some unicellular species. *Second*, monophyly of the family Goniaceae is not supported. *Third*, section *Volvox* is inferred to be sister to the remaining Volvocaceae. *Fourth*, cellular differentiation independently arose at least four and perhaps as many as six times within the volvocine algae.

### Colonial volvocine algae are not monophyletic

All three of our phylogenetic analyses indicate that the colonial volvocines are not monophyletic (**Figures 2** and **3**); further, an <u>A</u>pproximately <u>U</u>nbiased (AU) test strongly rejected monophyly for this group (p=2.82e-38) (**Supplementary Materials: Figure S1A**). These findings represent a major departure from earlier chloroplast gene-based volvocine phylogenies (Hanschen *et al*., 2018; Herron *et al*., 2009; Herron & Michod, 2008; Hu et al., 2019; Nakada *et al*., 2010; Nakazawa *et al*., 2001; Nozaki *et al*., 2000, 2014), phylogenies based on morphological characters (Nozaki, 1996; Nozaki & Itoh, 1994), phylogenies inferred using ITS 1 and 2 sequences (Coleman, 1999), as well as less taxonomically comprehensive phylogenies inferred using nuclear data (Zhang *et al*., 2019), all of which suggest that the colonial volvocine algae are monophyletic.

**Figure 2.**
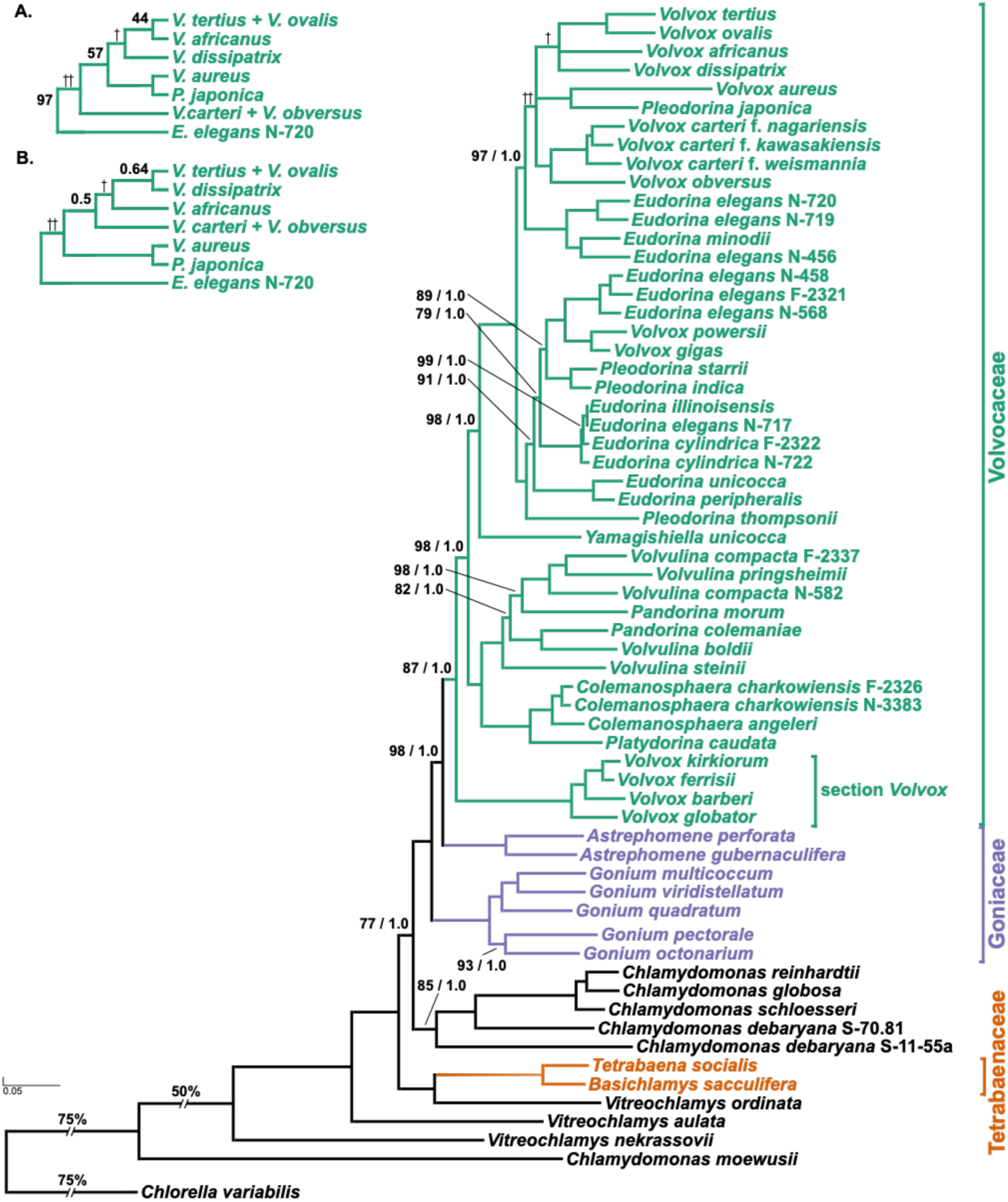
Molecular phylogeny of the colonial volvocine algae (Tetrabaenaceae, Goniaceae, and Volvocaceae) and closely related unicellular taxa represented by *Chlamydomonas* and *Vitreochlamys* with *Chlorella variabilis* as the outgroup. The phylogenetic tree shown is based on a multi-gene dataset of single-copy, protein-coding nuclear genes (12,650 aligned amino acid positions of 68 taxa) inferred using the maximum likelihood method, the branching order of which is nearly identical to that inferred in the Bayesian Inference using MrBayes. Numbers on branches represent bootstrap values and Bayesian posterior probabilities, respectively (all support values not shown are MLBS = 100, BPP = 1.0). Branch lengths correspond to genetic divergence, as indicated by the scale bar. Members of the Tetrabaenaceae, Goniaceae, and Volvocaceae are denoted in orange, purple, and green, respectively; unicellular species are denoted in black. Daggers correspond to branching order differences between the maximum likelihood and Bayesian inference analyses that are represented in greater detail in Figures 2A and 2B, respectively.

**Figure 3.**
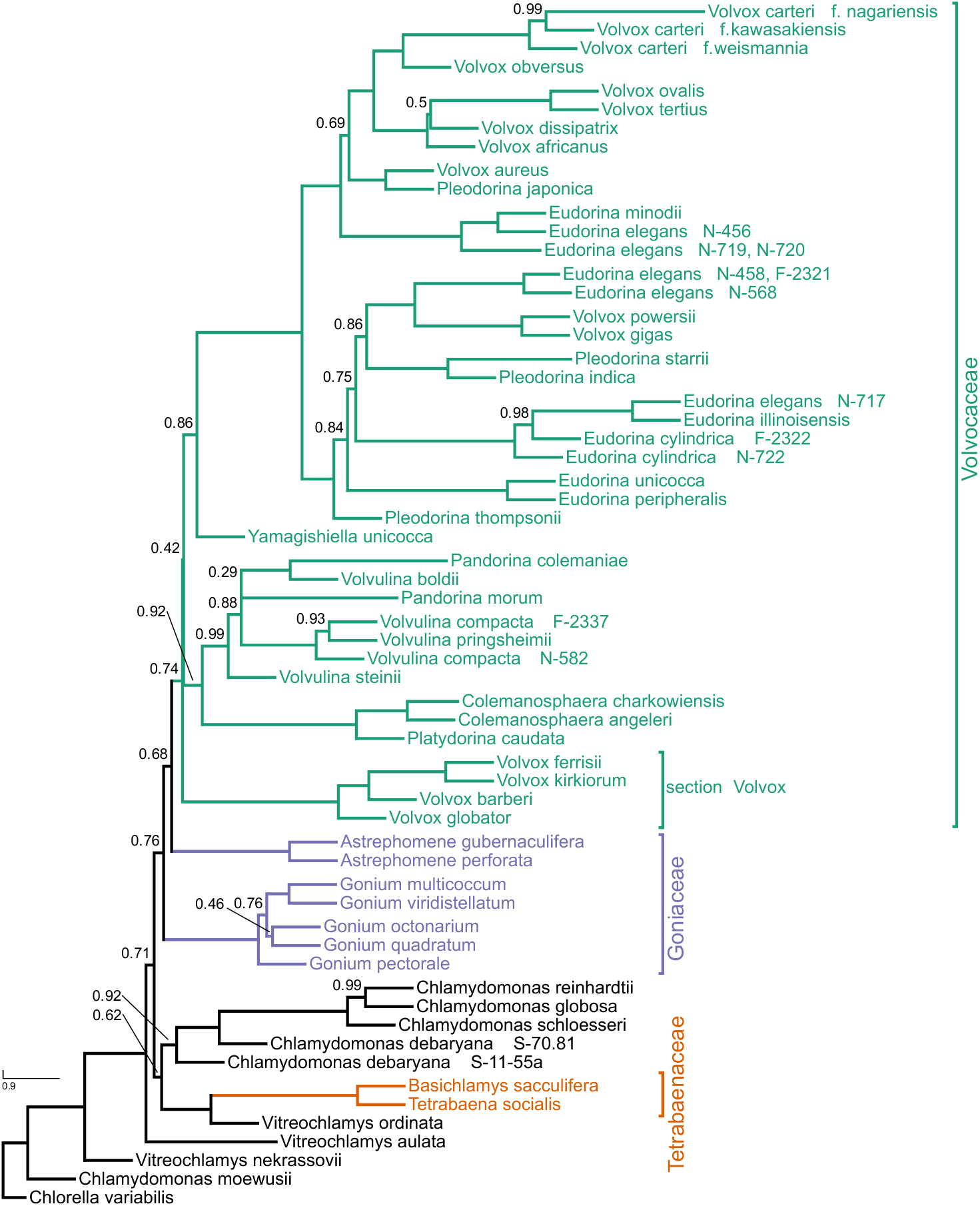
Phylogeny of the volvocine algae inferred using a coalescent-based analysis of 40 single-gene phylogenies. Numbers on branches represent posterior probabilities (support values not shown are CPP = 1.0). Members of the Tetrabaenaceae, Goniaceae, and Volvocaceae are denoted in orange, purple, and green, respectively; unicellular species are denoted in black.

Consistent with Pröschold *et al*. (2018), our results support the view that multicellularity evolved independently in the Tetrabaenaceae and in the Goniaceae + Volvocaceae. In each analytical framework, the Tetrabaenaceae was found to be sister to *Vitreochlamys ordinata* rather than to the Goniaceae + Volvocaceae (<u>M</u>aximum <u>L</u>ikelihood <u>B</u>oot<u>s</u>trap [MLBS] =100, <u>B</u>ayesian <u>P</u>osterior <u>P</u>robabilities [BPP]=1.0, Coalescent Posterior Probabilities [CPP]=1.0). A sister relationship between the Tetrabaenaceae and *V. ordinata* was inferred in 15/39 of our single-gene phylogenies and in 29/39 of our 4-taxa, unrooted, single-gene phylogenies (**Figure 4**). However, the maximum likelihood and Bayesian inference analyses differ from the coalescent-based analysis in the phylogenetic position of the *Chlamydomonas* clade. In MLBS, Tetrabaenaceae + *V. ordinata* is sister to a clade containing several species of *Chlamydomonas* (including *C. reinhardtii*), the Goniaceae, and the Volvocaceae (MLBS=100, BPP=1.0) (**Figure 2**). In the coalescent-based analysis, several species of *Chlamydomonas* are sister to the Tetrabaenaceae +*V. ordinata* (CPP=0.62) in a clade that is itself sister to the Goniaceae + Volvocaceae (CPP=0.71) (**Figure 3**). Although the branching order of *C. reinhardtii*, Tetrabaenaceae + *V. ordinata*, and Goniaceae + Volvocaceae differs between analyses, our results overall imply one independent origin of multicellularity in the Tetrabaenaceae and another origin in the Goniaceae + Volvocaceae.

**Figure 4.**
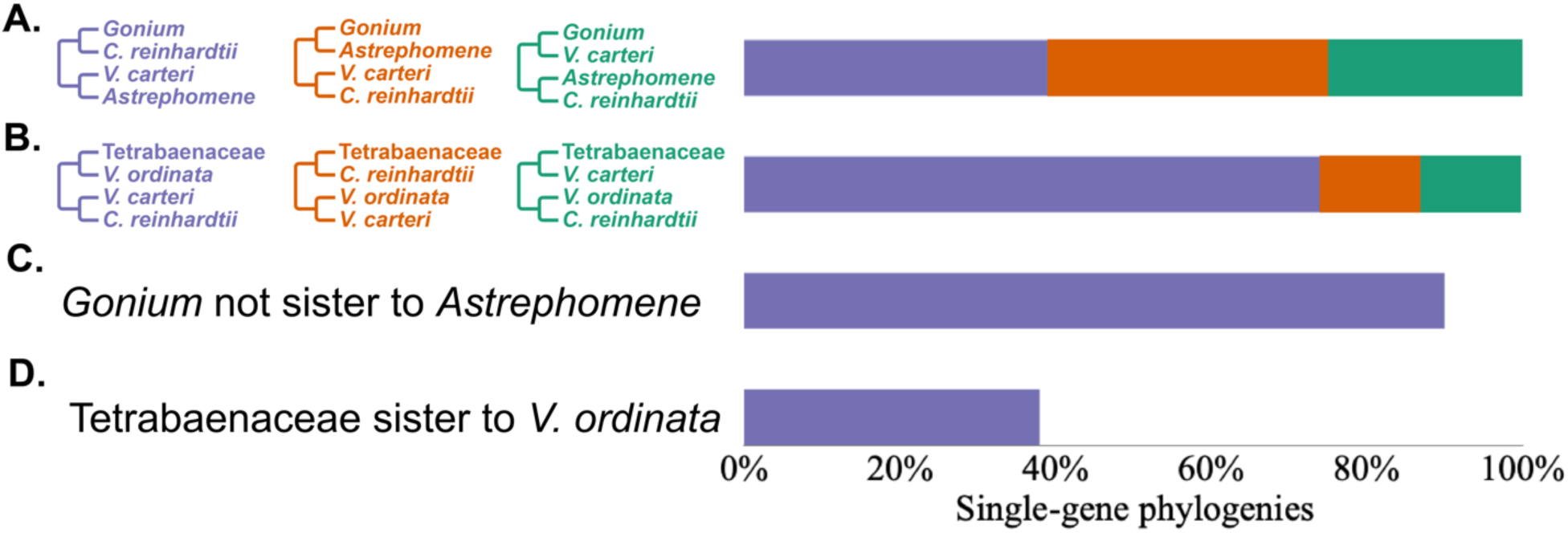
Phylogenetic relationships between *Gonium* and *Astrephomene* and between the Tetrabaenaceae and *Vitreochlamys ordinata*. Four-taxon, unrooted trees were generated by collapsing our single-gene phylogenies. The percentage of single-gene phylogenies representing a specific four-taxon, unrooted tree is represented by the purple, orange, and green bars. **(A)** percentage of four-taxon, unrooted trees representing specific relationships between *Gonium* and *Astrephomene*. **(B)** percentage of four-taxon, unrooted trees representing specific relationships between the Tetrabaenaceae and *V ordinata* in four-taxon, unrooted trees. **(C)** percentage of single-gene phylogenies that show *Gonium* not sister to *Astrephomene* represented by the purple bar. **(D)** percentage of single-gene phylogenies that show Tetrabaenaceae sister to *V. ordinata* represented by the purple bar. For all relationships involving *V. ordinata*, 39 out of 40 single-gene phylogenies were used due to *V. ordinata* not appearing in one of the inferences. All single-gene phylogenies were inferred using maximum likelihood under the appropriate evolutionary model as estimated by ProtTest.

Our results differ in key respects from a recent volvocine algae phylogeny offered by Zhang *et al*. (2019), which like ours is based on single-copy nuclear genes. Zhang *et al*. (2019) sought to understand the evolutionary relationships between two psychrophilic algae: *Chlamydomonas* sp. ICE-L and *Tetrabaena socialis* N-691. To do so they constructed a phylogeny consisting of ICE-L, N-691, three colonial *Volvox* strains, and eight unicellular species, including *C. reinhardtii*. Among their conclusions was that *T. socialis* N-691 is sister to the Volvocaceae, which is at odds with results shown in **Figures 2** and **3**. These results indicate that the Tetrabaenaceae is sister to *V. ordinata*, and together they are sister to *C. reinhardtii* + Goniaceae + Volvocaceae.

We hypothesized that the lack of concordance between our findings and those of Zhang *et al*. (2019) could be attributed to limited taxon sampling. To test this hypothesis, we first confirmed that *T. socialis* N-691 and *T. socialis* N-571 are conspecific. Nozaki and Ohtani (1992) examined *T. socialis* N-691 and found that N-691 was morphologically identical to *T. socialis* N-571 except for forming slightly larger colonies. Next, we compared ITS-1, 5.8S rRNA, and ITS-2 sequences from Mai & Coleman (1997) between the two *T. socialis* strains; we found that they were 92% identical. Then, we performed a BLAST search of the aforementioned sequences: the first ten BLAST hits all matched the same sequences from *T. socialis* strains with 100% query coverage. Using the BLAST software, neighbor joining and fast minimum evolution distance trees were inferred from the BLAST results; the trees produced show *T. socialis* strains N-571 and N-691 to be reciprocally monophyletic. Once we confirmed that N-691 and N-571 were conspecific, we were able to replicate the branching order produced by Zhang *et al*. (2019) using our concatenated 40-gene dataset (**Supplementary Materials: Figure S2A**). For our initial tree, we sampled our strains of *Chlamydomonas reinhardtii*, *C. moewusii*, *T. socialis*, *Volvox aureus*, *V. carteri* f. *nagariensis*, and *V. globator* to match taxa that were used in that study. For an outgroup species we sampled *Chlorella variabilis*. Multiple studies have shown that the accuracy of phylogenetic reconstruction can be improved by increasing the number of taxa sampled (Hedtke et al., 2006; Pollock et al., 2002; Zwickl & Hillis, 2002). When we added more taxa and performed Maximum Likelihood analysis on the new dataset, the three colonial volvocine families were no longer monophyletic. The Tetrabaenaceae were sister to *Vitreochlamys ordinata*, and this clade appeared sister to *C. reinhardtii* + Goniaceae + Volvocaceae. (**Supplementary Materials: Figure S2B**). These analyses confirm that the placement of *T. socialis* N-691 as sister to the Volvocaceae is an artefact of limited taxon sampling. From this we draw three conclusions: *First*, the colonial volvocine algae are not monophyletic; s*econd,* at least two independent origins of multicellularity occurred within the volvocine algae; *third,* once multicellularity evolved no extant lineage reverted to the ancestral unicellular state (see **Figures 2** and **3**).

### The family Goniaceae is not monophyletic

Multiple volvocine phylogenies have concluded that the Goniaceae is monophyletic (Hanschen *et al*., 2018a, 2018b; Herron *et al*., 2009; Herron & Michod, 2008; Nakada *et al*., 2019; Nozaki, 1996; Nozaki *et al*., 1996, 1997, 2000; Nozaki & Itoh, 1994; Pröschold *et al*., 2018). Our analyses suggest otherwise (**Figure 2** and **3**): we find that *Astrephomene* is sister to the Volvocaceae (MLBS= 98, BPP= 1.0, CBPP= 0.68) rather than to *Gonium*. This inference is strengthened by observations that 36/40 of our single-gene phylogenies show that *Gonium* and *Astrephomene* are not sister taxa, as do 23/40 of our four-taxon, unrooted phylogenies (**Figure 4**). All three of our analyses indicate that *Astrephomene* is monophyletic and sister to the Volvocaceae clade (MLBS=98, BPP=1.0, CPP=0.68), with *Gonium* sister to *Astrephomene* + Volvocaceae (MLBS=100, BPP=1.0, CPP=0.76). Furthermore, we performed an AU test where the monophyly of the Goniaceae was tested against our finding of paraphyly for the Goniaceae. The null hypothesis, monophyly of the Goniaceae, was rejected (p=0.0446) (**Supplementary Materials: Figure S1B**).

Prior studies have produced mixed results regarding monophyly of the Goniaceae, sometimes with low support values for the relevant relationships. Nozaki and colleagues (1995) published four phylogenies inferred using a single chloroplast gene and different inference methods; all four trees either showed low support for monophyly of the Goniaceae or suggested a topology where *Astrephomene* is sister to *Gonium* + Volvocaceae. Coleman (1999) inferred a volvocine phylogeny based on ITS-1 and ITS-2 sequences that showed *Astrephomene* sister to Tetrabaenaceae *+ Gonium* + Volvocaceae; however, the bootstrap support for this suggested relationship was between 50% and 75%, indicating weak support for the branching order. Other phylogenies suggesting monophyly in the Goniaceae do so with weak or contradictory support (Nakada *et al*., 2019; Nozaki *et al*., 2000; Pröschold *et al*., 2018).

Our inference that Goniaceae are not monophyletic is consistent with some – but not all – of the analyses recently reported by Pröschold *et al*. (2018) and Nakada *et al*. (2019). However, we should not disregard past morphological and ultrastructural studies suggesting a close relationship between *Astrephomene* and *Gonium* (Nozaki, 1990; Nozaki & Itoh, 1994; Nozaki & Kuroiwa, 1992). These taxa differ from the Volvocaceae in that each cell, rather than the entire colony, is surrounded by a tripartite boundary (Kirk *et al*., 1986). This feature distinguishes their mode of colony formation from all other colonial algae within the Volvocaceae; our results suggest that it is ancestral to the Goniaceae + Volvocaceae and lost in the Volvocaceae.

### Volvox section Volvox is sister to the remaining Volvocaceae

Our data indicate that *Volvox* section *Volvox* is not a subclade within either the *Pandorina* + *Volvulina* + *Colemanosphaera* (PVC) or *Eudorina + Volvox +Pleodorina* (EVP) subclades. Older studies based on the *rbcL* chloroplast gene (Nozaki *et al*., 1996), ITS-1 and ITS-2 sequences (Coleman, 1999), and morphology (Nozaki and Itoh, 1994) suggest that section *Volvox* belongs to a clade that encompasses *Eudorina*, *Pleodorina*, and other *Volvox* species. More recent studies of the volvocine algae based on 5 chloroplast genes, or based on multiple datasets that include 1 chloroplast gene (Pröschold *et al*. (2018), suggest that section *Volvox* belongs to a clade that includes *Pandorina*, *Volvulina*, *Platydorina* (Hanschen *et al*., 2018a; Herron & Michod, 2008), and (in the studies where it was included) *Colemanosphaera* (Hu *et al*., 2019; Nozaki *et al*., 2014). By contrast, all of our analyses indicate that section *Volvox* is monophyletic and sister to the remaining Volvocaceae (MLBS=87, BPP=1.0, CPP= 0.74). AU tests rejected both section *Volvox* being sister to *Colemanosphaera* + *Platydorina* (p-AU=4.64e-88) or sister to the PVC clade (p-AU=0.0332) (**Supplementary Materials: Figure S1C**). These results bolster our finding that section *Volvox* is sister to the remaining Volvocaceae (**Figures 2** and **3**).

### Cellular differentiation independently arose at least four times in the volvocine algae

The last major difference between our results and earlier phylogenies concerns the number of independent origins of cellular differentiation. Prior literature suggests that cellular differentiation independently evolved at least three times: once in *Astrephomene*, once in section *Volvox,* and at least once in the EVP clade (Grochau-Wright *et al*., 2017; Herron & Michod, 2008). By contrast, our results show a *minimum* of four independent origins of cellular differentiation: one in *Astrephomene*, one in section *Volvox*, and at least two in the EVP clade (**Figure 5A**). We cannot exclude the possibility of two additional independent origins in the branches leading to *Pleodorina starrii* and *Volvox gigas* (**Figure 5A**). In *Astrephomene*, section *Volvox, Pleodorina*, and *Volvox dissipatrix*, differentiated cells carry out the function of motility, whereas undifferentiated cells participate in both motility and reproduction (Kirk, 2005). The remaining *Volvox* species within the EVP clade have all evolved specialized germ cells for reproduction and somatic cells for motility (Herron *et al*., 2009, 2010).

**Figure 5.**
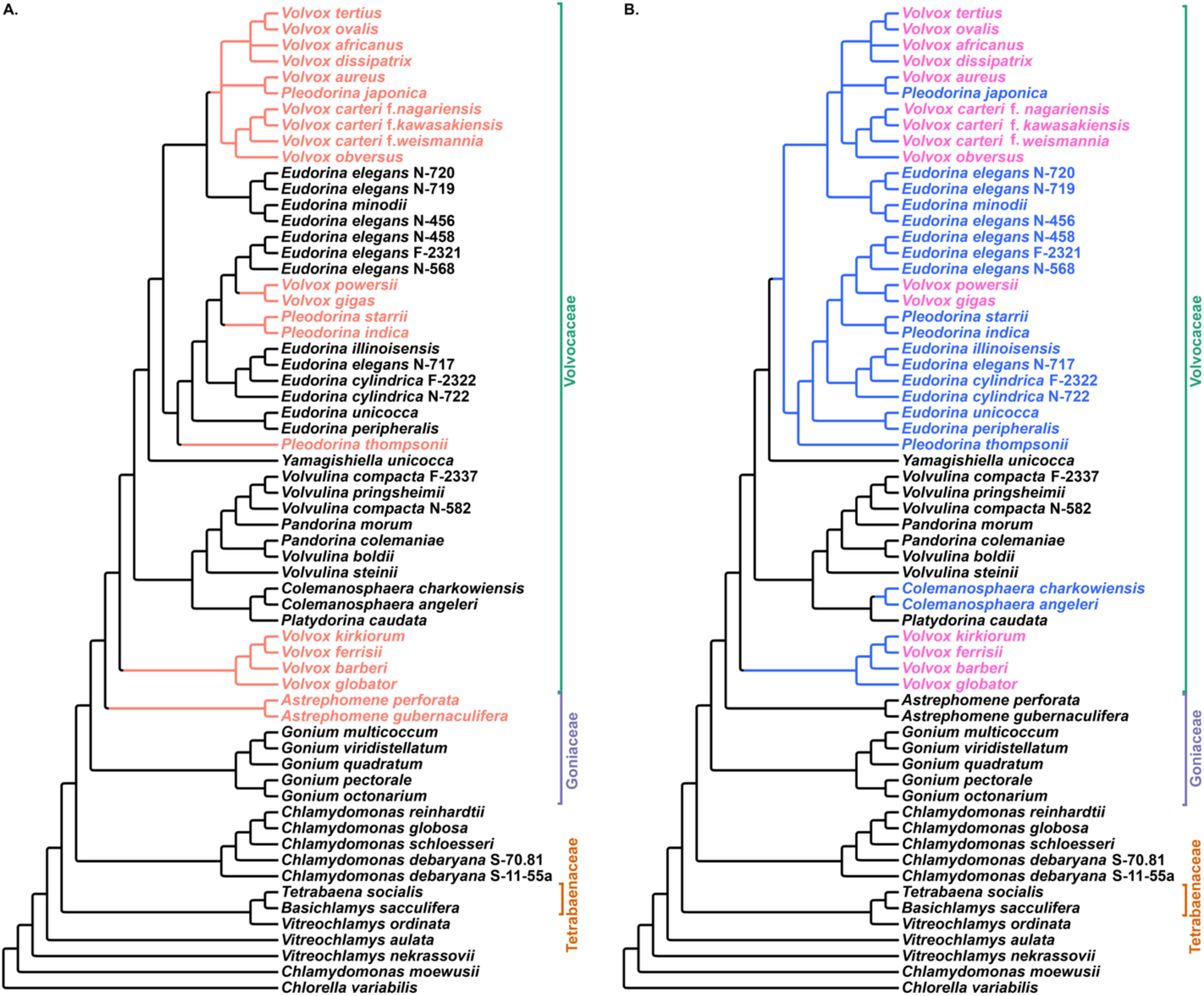
**(A)** Phylogeny of the volvocine algae highlighting the lineages in which soma differentiation has evolved (peach). This tree indicates a minimum of four and maximum of six independent origins of cellular differentiation **(B)** Phylogeny of the volvocine algae highlighting the lineages that are isogamous (black), anisogamous (blue), and oogamous (names in pink only). Both phylogenies were inferred using maximum likelihood.

### Isogamy is the ancestral mode of sexual reproduction

Consistent with past studies, our results suggest that isogamy, the production of similar sized, motile gametes, is the ancestral mode of sexual reproduction among the volvocine algae (**Figure 5B** and **Supplementary Materials: Table S2**). Isogamy is present in the unicellular genera *Chlamydomonas* and *Vitreochlamys*, and is retained within the multicellular genera *Astrephomene*, *Basichlamys*, *Gonium*, *Pandorina*, *Platydorina*, *Tetrabaena*, *Volvulina*, and *Yamagishiella*. *Colemanosphaera*, *Eudorina*, *Pleodorina*, and *Volvox* have all evolved either anisogamy or oogamy (Herron, 2016; Nozaki *et al*., 2014; da Silva, 2018; Umen, 2011). Anisogamy appears to have independently evolved at least three times from an isogamous ancestor: in section *Volvox* and in both *Colemanosphaera* and EVP. Conventional anisogamy, which consists of two motile gamete types of unequal size, appears in *Colemanosphaera*, *Eudorina*, and *Pleodorina.* Oogamy, a specialized form of anisogamy where the female gamete is immotile and significantly larger than the motile, male gamete, is inferred to have independently evolved at least three times in lineages leading to section *Volvox*, *V. gigas* + *V. powersii*, and in the clade containing *V. africanus*, *V. aureus*, *V. carteri*, *V. dissipatrix*, *V. obversus*, *V. ovalis*, and *V. tertius* (Herron, 2016; Nozaki *et al*., 2014; da Silva, 2018). This latter finding differs from Hanschen *et al*. (2018b) who reported at least two independent origins of oogamy among the volvocine algae. Our results clearly suggest an independent origin of oogamy in section *Volvox*. Due to the branching order of the Hanschen *et al*. (2018b) phylogeny, however, we cannot be sure that the most recent common ancestor of *V. gigas* and *V. powersii* and *V. carteri* and other *Volvox* species within that clade was not oogamous.

### Platydorina caudata is sister to Colemanosphaera, and Pandorina is paraphyletic with respect to Volvulina

Within the PVC clade, our results add further support to the view that *Pandorina* is paraphyletic with respect to *Volvulina* (MLBS=100, BPP=1.0, CPP=0.92) (Coleman, 1999, 2001; Hanschen *et al*., 2018b; Herron *et al*., 2009; Herron & Michod, 2008; Nozaki *et al*., 2000, 2014). Also, consistent with other multi-gene analyses *Colemanosphaera* appears to be monophyletic with high support (MLBS=100, BPP=1.0, CPP=1.0) and sister to *Platydorina* (MLBS=100, BPP=1.0, CPP=1.0) (**Figures 2** and **3**) (Hanschen *et al*., 2018a; Nozaki *et al*., 2014, 2015).

### The genera Eudorina, Volvox and Pleodorina are polyphyletic

*Yamagishiella unicocca* is sister to the *<u>E</u>udorina+<u>V</u>olvox+Pleodorina* (EVP) clade, which encompasses two large subclades (MLBS=98, BPP=1.0, CPP=0.86) (**Figures 2** and **3**). Our results support prior work suggesting that the genera *Volvox*, *Eudorina* and *Pleodorina* are not monophyletic (Coleman, 1999; Grochau-Wright *et al*., 2017; Hanschen *et al*., 2018a, 2018b; Herron *et al*., 2009, 2010; Herron & Michod, 2008; Nozaki, 2003; Nozaki *et al*., 2000, 2006, 2014, 2015). The genus *Volvox* appears to be polyphyletic, with members represented across the two EVP subclades and the section *Volvox* clade. Members of both the *Pleodorina* and *Eudorina* genera are inferred to be polyphyletic across the two EVP subclades.

Historically, the genus *Volvox* has been divided into 4 sections – *Copelandosphaera*, *Janetosphaera*, *Merrillosphaera*, and *Volvox* – based on morphological (Smith, 1944) and molecular data (Nozaki, 2003). A recent section-level revision of the genus *Volvox* (Nozaki *et al*., 2015) resulted in the creation and deletion of sections *Besseyosphaera* and *Copelandosphaera*, respectively. Hereafter, we will only refer to the revised taxonomic sections proposed by Nozaki *et al*. (2015), with which our Bayesian inference and coalescent-based results are in agreement (**Supplementary Materials: Figure S3**). Our Bayesian inference and coalescent-based analyses further suggest that each of the four sections is monophyletic, and that none encompass novel taxa not listed by Nozaki *et al*. (2015). The branching order of our ML analysis, however, suggests that section *Merrillosphaera* is not monophyletic (**Supplementary Materials: Figure S3A**), although the support values for the relevant relationships are low. Specifically, our ML analysis suggests that *V. africanus*, *V. dissipatrix*, *V. ovalis*, and *V. tertius* form a clade with *V. aureus* and *P. japonica* that is separate from the other *Merrillosphaera* taxa (MLBS=59). In contrast, our BI and CB analyses suggest with moderate support (BPP=0.50, CPP=0.69) that the *Merrillosphaera* species are monophyletic (**Supplementary Materials: Figure S3B**). Heeding our support values rather than only the branching order, we propose that the taxonomic system of the genus *Volvox* as outlined by Nozaki and colleagues (2014) be retained.

### Unicellular taxa are nested within the clade containing the colonial volvocine algae

Of the unicellular taxa, *Chlamydomonas debaryana, C. globosa*, *C. reinhardtii*, *C. schloesseri*, and *Vitreochlamys ordinata* are nested within the clade containing the colonial volvocine algae. Our results confirm prior studies showing the genus *Vitreochlamys* to be polyphyletic (Nakada *et al*., 2019; Nakazawa *et al*., 2001). The closest unicellular relative to the clade that contains the colonial algae + *C. reinhardtii* is suggested to be *V. aulata* (**Figures 2** and **3**). This suggests that at least some members of *Vitreochlamys* are very closely related to the colonial volvocine algae. This relationship had been previously suggested by other studies (Nakada *et al*., 2019; Nozaki *et al*., 1999) including Nakazawa *et al*. (2001), whose ultrastructural studies uncovered striking similarities in how these taxa formed pyrenoids and eyespot apparati (stigma), and established their tripartite cell walls.

*Chlamydomonas* is a polyphyletic genus (Craig *et al*., 2021; Nakada *et al*., 2008, 2019; Pröschold *et al*., 2001) composed of at least 500 species (Pröschold *et al*., 2001). Although we sampled only a handful of *Chlamydomonas* species, our data support this view and broadly agree with the *Chlamydomonas* relationships inferred by Pröschold and colleagues (2018), who used a combination of molecular phylogenetic analyses, sporangium wall lysis tests, and ultrastructural analyses. Our data strongly support *C. schloesseri* being sister to *C. reinhardtii* + *C. globosa* (MLBS=100, BPP=1.0, CPP=1.0) and designating *C. schloesseri* as a “true” *Chlamydomonas* species, as suggested by Pröschold *et al*. (2018). Our study is also in agreement with a recent study by Craig *et al*. (2021) that shows *C. schloesseri* being sister to *C. reinhardtii* + *C. globosa*. Also, like Pröschold *et al*. (2018), our analyses indicate that *C. debaryana* SAG 70.81 is sister to *Chlamydomonas schloesseri* and its relatives (MLBS=100, BPP= 1.0, CPP=1.0). However, unlike the Pröschold *et al*. (2018) study which proposed that strain *C. debaryana*/*Edaphochlamys debaryana* (SAG 11-55a) is sister to the Tetrabaenaceae, our analyses support the view that *C. debaryana*/*Edaphochlamys debaryana* is more closely related to *C. reinhardtii* (MLBS=84, BPP=1.0, CPP=0.92) than to the colonial algae. Our finding is further supported by Craig *et al*. (2021) who inferred that *C. debaryana*/*Edaphochlamys debaryana + Chlamydomonas sphaeroides* is sister to the clade containing *C. schloesseri* + *C. reinhardtii* + *C. globosa*. Our placement of *C. debaryana* (SAG 11-55a) could be a result of limited (N=6) sampling within the *Chlamydomonas* genus, which was more extensively sampled by Pröschold *et al*. (2018) (N>30). Consistent with a prior study, *C. moewusii* appears to be more distantly related to the colonial volvocines than is *Vitreochlamys nekrassovii* (Herron & Michod, 2008).

## Conclusions

Using a 40-protein dataset, we have shown that the Tetrabaenaceae and the Goniaceae + Volvocaceae likely represent two independent origins of multicellularity and that cellular differentiation has independently evolved at least four, and possibly six, times within the volvocine algae. Our results suggest that both multicellularity and cellular differentiation are evolutionary labile traits within the volvocine algae. We have established a robust phylogeny of this group, which we hope will assist future efforts aimed at re-evaluating ancestral character states and understanding the origins of multicellularity and cellular differentiation in the volvocine green algae. Future research directions that can be taken using our transcriptomic dataset include ancestral-state reconstruction of traits involving cellularity, cellular differentiation, and gamete size as well as discerning the evolutionary history of gene families across the volvocine algae as a whole and within its major clades.

## Data Archiving

All raw data generated and used for this study have been deposited in the National Center for Biotechnology Information (NCBI) Sequence Read Archive (SRA) under BioProject PRJNA701495. Accession numbers for our raw RNA-Seq reads range from SAMN17884613 to SAMN17884667. For detailed information regarding accession number assignment to a specific taxon please refer to **Table S5** in **Supplementary Materials.**

## Supporting information

Supplemental RNA Extraction Protocol Methods

translate_scaffold.txt

## Notes

### Competing Interest Statement

The authors have declared no competing interest.

### Summary of Updates

This version of the manuscript has been revised to update the Results and Discussion sections due to concerns regarding length. This version has a combined Results and Discussion section which shortened and improved the readability of the manuscript.

