## Supplemental Tables and Figures for "Phylotranscriptomics points to multiple independent origins of multicellularity and cellular differentiation in the volvocine algae"

**Supplementary Table S1.** Medium used to culture each sequenced strain. Under Collection ID, "NIES" refers to National Institute for Environmental Studies (NIES, Japan), "SAG" refers to Culture Collection of Algae at the University of Göttingen (SAG, Germany), and "UTEX" refers to The Culture Collection of Algae at the University of Texas at Austin (UTEX, USA). AF-6 (Kato, 1982), AF-6/2 (Kato, 1982), CA (Ichimura & Watanabe, 1974), MG (Ichimura, 1973), and VT (Starr, 1973) were prepared according to media recipes supplied by NIES. HS media (Sueoka, 1960) was prepared according to media recipe supplied by Chlamydomonas Resource Center. Soilwater: GR+ media was purchased from UTEX.

| **Taxa** | **Collection ID** | **Culture Medium** |
|---|---|---|
| *Astrephomene gubernaculifera* | NIES-418 | VT |
| *A. perforata* | NIES-564 | VT |
| *Basichlamys sacculifera* | NIES-566 | AF-6 |
| *Colemanosphaera charkowiensis* | NIES-3383 | AF-6 |
| *Chlamydomonas debaryana* | SAG 11-55a | HS |
| | SAG 70.81 | HS |
| *C. globosa* | SAG 81.72 | HS |
| *C. moewusii* | SAG 11-16f | HS |
| *C. schloesseri* | SAG 2486 | HS |
| *Eudorina cylindrica* | NIES-722 | AF-6 |
| *E. elegans* | NIES-456 | VT |
| | NIES-458 | VT |
| | NIES-568 | CA |
| | NIES-717 | AF-6 |
| | NIES-719 | VT |
| | NIES-720 | VT |
| *E. illinoisensis* | NIES-460 | VT |
| *E. minodii* | NIES-856 | AF-6 |
| *E. peripheralis* | NIES-725 | AF-6 |
| *E. unicocca* | SAG 24-1c | AF-6 |
| *Gonium multicoccum* | NIES-737 | VT |
| *G. octonarium* | NIES-851 | AF-6 |
| *G. quadratum* | NIES-653 | AF-6 |
| *G. viridistellatum* | NIES-654 | VT |
| *Pandorina colemaniae* | NIES-572 | AF-6 |
| *P. morum* | NIES-890 | AF-6 |
| *Platydorina caudata* | NIES-728 | MG |
| *Pleodorina indica* | NIES-736 | AF-6 |
| *P. japonica* | UTEX 2523 | Soilwater: GR+ |
| *P. starrii* | NIES-1362 | AF-6 |
| | NIES-1363 | AF-6 |
| *P. thompsonii* | NIES-4126 | AF-6 |
| *Vitreochlamys aulata* | NIES-878 | AF-6 |
| | SAG 80.81 | AF-6 |
| *V. nekrassovii* | SAG 11-10 | AF-6 |

| | | |
|---|---|---|
| *V. ordinata* | NIES-882 | AF-6 |
| *Volvox africanus* | NIES-863 | AF-6/2 |
| *V. aureus* | NIES-541 | VT |
| *V. barberi* | NIES-730 | AF-6 |
| *V. carteri* f. *kawasakiensis* | NIES-732 | VT |
| *V. carteri* f. *nagariensis* | NIES-865 | MG |
| *V. carteri* f. *weismannia* | NIES-866 | VT |
| *V. dissipatrix* | NIES-4128 | AF-6 |
| *V. ferrisii* | NIES-3986 | AF-6 |
| *V. gigas* | NIES-867 | MG |
| *V. globator* | SAG 199.80 | AF-6 |
| *V. kirkiorum* | NIES-543 | VT |
| *V. obversus* | NIES-868 | AF-6 |
| *V. ovalis* | NIES-2569 | AF-6 |
| *V. powersii* | NIES-4127 | AF-6 |
| *V. tertius* | NIES-544 | AF-6 |
| *Volvulina boldii* | NIES-893 | MG |
| *V. compacta* | NIES-582 | VT |
| *V. pringsheimii* | NIES-895 | MG |
| *V. steinii* | SAG 90-1 | AF-6 |

**Supplementary Table S2.** Information on sampled genera regarding cellularity, typical cell number, differentiation, and gamete size.

| | Typical Cell # | Differentiation | Gamete Size |
|---|---|---|---|
| **Unicellular Genera** | | | |
| *Chlorella* | 1 | - | Isogamy |
| *Chlamydomonas* | 1 | - | Isogamy |
| *Vitreochlamys* | 1 | - | Isogamy |
| **Multicellular Genera** | | | |
| *Astrephomene* | 32-128 | Somatic | Isogamy |
| *Basichlamys* | 4 | - | Isogamy |
| *Colemanosphaera* | 16-32 | - | Anisogamy |
| *Eudorina* | 16-32 | - | Anisogamy |
| *Gonium* | 8-32 | - | Isogamy |
| *Pandorina* | 16-32 | - | Isogamy |
| *Platydorina* | 16-32 | - | Isogamy |
| *Pleodorina* | 32-128 | Somatic | Anisogamy |
| *Tetrabaena* | 4 | - | Isogamy |
| *Volvox* | 500-50,000 | Somatic and Germ* | Oogamy |
| *Volvulina* | 8-16 | - | Isogamy |
| *Yamagishiella* | 16-32 | - | Isogamy |

* *Volvox* lineages that have evolved specialized somatic and germ cell lines are: *Volvox africanus*, *V. carteri*, *V. gigas*, and *V. obversus*. All other *Volvox* species have specialized somatic cells where undifferentiated cells partake in motility and reproductive functions.

**Supplementary Table S3.** Accession numbers of previously published RNA-seq data. Under Collection ID, “FACHB” refers to Culture Collection of Freshwater Algae at the Institute of Hydrobiology, Chinese Academy of Sciences (FACHB, China), and “NIES” refers to National Institute for Environmental Studies (NIES, Japan). All FACHB strains used in this study were sequenced by Hu et al. (2020) under BioProject PRJNA532307. *Tetrabaena socialis* NIES-571 was sequenced by Featherston et al. (2018) under BioProject PRJNA393411.

| RNA-seq Data Used | Collection ID | Accession # |
|---|---|---|
| *Colemanosphaera angeleri* | FACHB 2363 | SRX5666821 |
| *C. charkowiensis* | FACHB 2326 | SRX5666822 |
| *Eudorina cylindrica* | FACHB 2322 | SRX5666825 |
| *E. elegans* | FACHB 2321 | SRX5666826 |
| *Pandorina colemaniae* | FACHB 2361 | SRX5666819 |
| *P. morum* | FACHB 2362 | SRX5666823 |
| *Tetrabaena socialis* | NIES-571 | SRX3367144 |
| *Volvulina compacta* | FACHB 2337 | SRX5666820 |
| *Yamagishiella unicocca* | FACHB 2364 | SRX5666824 |

**Supplementary Table S4.** 40 genes with best predicted evolutionary model under the Akaike information criterion (AIC). Gene names from De Clerck et al., (2018) were used and unchanged. Under Evolutionary model, “I” refers to invariable sites, “G” refers to gamma-distributed rates, “F” refers to empirical frequency estimation, “LG” refers to the Le & Gascuel substitution model (2008), “JTT” refers to the Jones, Taylor, and Thornton substitution model (Jones et al., 1992), “WAG” refers to the Whelan and Goldman substitution model (Whelan & Goldman, 2001), and “DAYHOFF” refers to the Dayhoff substitution model (Dayhoff et al., 1978).

| **Gene Name** | **Evolutionary model (AIC)** |
|---|---|
| HOM04ULVA001181 | LG+I+G+F |
| HOM04ULVA001650 | LG+I+G+F |
| HOM04ULVA001976 | JTT+I+G |
| HOM04ULVA002177 | JTT+I+G |
| HOM04ULVA002242 | JTT+I+G |
| HOM04ULVA002252 | LG+I+G |
| HOM04ULVA002273 | DAYHOFF+I+G |
| HOM04ULVA002313 | LG+I+G |
| HOM04ULVA002345 | JTT+I+G+F |
| HOM04ULVA002356 | LG+G+F |
| HOM04ULVA002369 | LG+I+G |
| HOM04ULVA002371 | LG+I+G+F |
| HOM04ULVA002373 | LG+I+G |
| HOM04ULVA002381 | LG+I+G |
| HOM04ULVA002396 | LG+I+G |
| HOM04ULVA002447 | LG+I+G |
| HOM04ULVA002537 | LG+G |
| HOM04ULVA002538 | JTT+G+F |
| HOM04ULVA002544 | LG+I+G+F |
| HOM04ULVA002581 | LG+G+F |
| HOM04ULVA002624 | JTT+I+G |
| HOM04ULVA002640 | LG+I+G |
| HOM04ULVA002681 | JTT+I+G |
| HOM04ULVA002762 | JTT+G |
| HOM04ULVA002763 | JTT+I+G+F |
| HOM04ULVA002810 | WAG+I+G |
| HOM04ULVA002814 | LG+I+G+F |
| HOM04ULVA002888 | JTT+G |
| HOM04ULVA002962 | JTT+I+G |
| HOM04ULVA002986 | LG+I+G+F |
| HOM04ULVA003057 | JTT+I+G |
| HOM04ULVA003095 | LG+G |
| HOM04ULVA003157 | JTT+I+G |
| HOM04ULVA003199 | JTT+I+G |
| HOM04ULVA003203 | JTT+I+G |
| HOM04ULVA003211 | LG+I+G+F |

| | |
|---|---|
| HOM04ULVA003266 | JTT+G |
| HOM04ULVA003755 | LG+I+G+F |
| HOM04ULVA003786 | LG+I+G+F |
| HOM04ULVA004274 | LG+I+G |

**Table S5.** National Center for Biotechnology Institute (NCBI) accession numbers for BioProject PRJNA701495. "NIES" refers to National Institute for Environmental Studies (NIES, Japan), "SAG" refers to Culture Collection of Algae at the University of Göttingen (SAG, Germany), and "UTEX" refers to The Culture Collection of Algae at the University of Texas at Austin (UTEX, USA).

| Taxa | Collection ID | NCBI Accession Number |
|---|---|---|
| *Astrephomene gubernaculifera* | NIES-418 | SAMN17884640 |
| *A. perforata* | NIES-564 | SAMN17884616 |
| *Basichlamys sacculifera* | NIES-566 | SAMN17884617 |
| *Colemanosphaera. charkowiensis* | NIES-3383 | SAMN17884629 |
| *Chlamydomonas debaryana* | SAG 11-55a | SAMN17884664 |
| | SAG 70.81 | SAMN17884633 |
| *C. globosa* | SAG 81.72 | SAMN17884666 |
| *C. moewusii* | SAG 11-16f | SAMN17884663 |
| *C. schloesseri* | SAG 2486 | SAMN17884667 |
| *Eudorina cylindrica* | NIES-722 | SAMN17884650 |
| *E. elegans* | NIES-456 | SAMN17884613 |
| | NIES-458 | SAMN17884614 |
| | NIES-568 | SAMN17884644 |
| | NIES-717 | SAMN17884647 |
| | NIES-719 | SAMN17884648 |
| | NIES-720 | SAMN17884649 |
| *E. illinoisensis* | NIES-460 | SAMN17884641 |
| *E. minodii* | NIES-856 | SAMN17884655 |
| *E. peripheralis* | NIES-725 | SAMN17884651 |
| *E. unicocca* | SAG 24-1c | SAMN17884665 |
| *Gonium multicoccum* | NIES-737 | SAMN17884621 |
| *G. octonarium* | NIES-851 | SAMN17884622 |
| *G. quadratum* | NIES-653 | SAMN17884646 |
| *G. viridistellatum* | NIES-654 | SAMN17884619 |
| *Pandorina colemaniae* | NIES-572 | SAMN17884618 |
| *P. morum* | NIES-890 | SAMN17884659 |
| *Platydorina caudata* | NIES-728 | SAMN17884652 |
| *Pleodorina indica* | NIES-736 | SAMN17884654 |
| *P. japonica* | UTEX 2523 | SAMN17884639 |
| *P. starrii* | NIES-1362 | SAMN17884626 |
| | NIES-1363 | SAMN17884627 |
| *P. thompsonii* | NIES-4126 | SAMN17884631 |
| *Vitreochlamys aulata* | NIES-878 | SAMN17884623 |
| | SAG 80.81 | SAMN17884634 |
| *V. nekrassovii* | SAG 11-10 | SAMN17884662 |
| *V. ordinata* | NIES-882 | SAMN17884624 |
| *Volvox africanus* | NIES-863 | SAMN17884637 |
| *V. aureus* | NIES-541 | SAMN17884642 |
| *V. barberi* | NIES-730 | SAMN17884653 |

| | | |
|---|---|---|
| *V. carteri* f. *kawasakiensis* | NIES-732 | SAMN17884620 |
| *V. carteri* f. *nagariensis* | NIES-865 | SAMN17884656 |
| *V. carteri* f. *weismannia* | NIES-866 | SAMN17884657 |
| *V. dissipatrix* | NIES-4128 | SAMN17884661 |
| *V. ferrisii* | NIES-3986 | SAMN17884630 |
| *V. gigas* | NIES-867 | SAMN17884658 |
| *V. globator* | SAG 199.80 | SAMN17884636 |
| *V. kirkiorum* | NIES-543 | SAMN17884643 |
| *V. obversus* | NIES-868 | SAMN17884638 |
| *V. ovalis* | NIES-2569 | SAMN17884628 |
| *V. powersii* | NIES-4127 | SAMN17884632 |
| *V. tertius* | NIES-544 | SAMN17884615 |
| *Volvulina boldii* | NIES-893 | SAMN17884660 |
| *V. compacta* | NIES-582 | SAMN17884645 |
| *V. pringsheimii* | NIES-895 | SAMN17884625 |
| *V. steinii* | SAG 90-1 | SAMN17884635 |

**Supplementary Figure S1.** Approximately Unbiased (AU) tests comparing key hypotheses for our 40-protein concatenated dataset. Each AU test used all 68 taxa from our study, but trees shown have been truncated so as to be easily viewed/read. T1 always represents the branching order of the unconstrained tree, and all other trees are constrained trees. Hypotheses tested: (A) Monophyly of the colonial volvocine algae, (B) Monophyly of Goniaceae, (C) Section *Volvox* sister to *Colemanosphaera* or PVC clade.

### A. Monophyly of the colonial volvocine algae

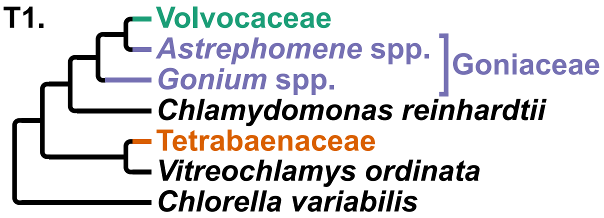


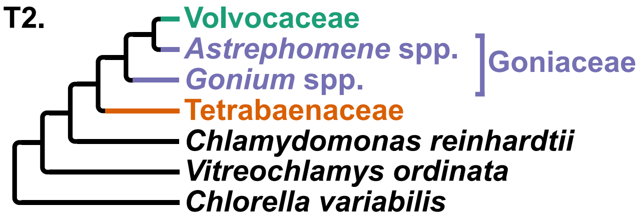


| Tree | logL | deltaL | p-AU |
|---|---|---|---|
| 1 | -308397.5847 | 0 | 1.0 + |
| 2 | -308653.5707 | 255.99 | 2.82e-38 - |

### B. Monophyly of Goniaceae

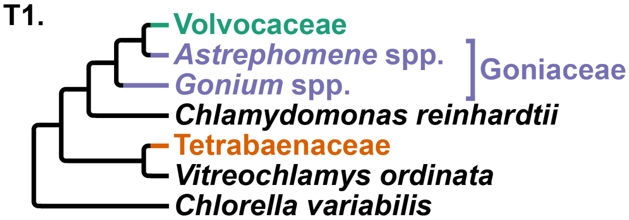


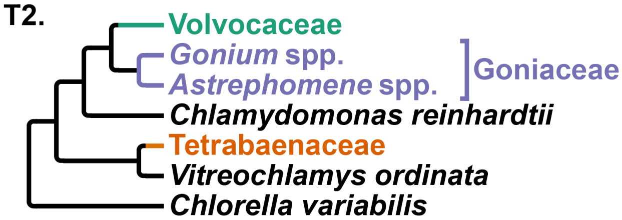


| Tree | logL | deltaL | p-AU |
|---|---|---|---|
| 1 | -308397.5785 | 0 | 0.955 + |
| 2 | -308429.6318 | 32.053 | 0.0446 - |

**C.** **Section *Volvox* sister to *Colemanosphaera* or PVC clade**

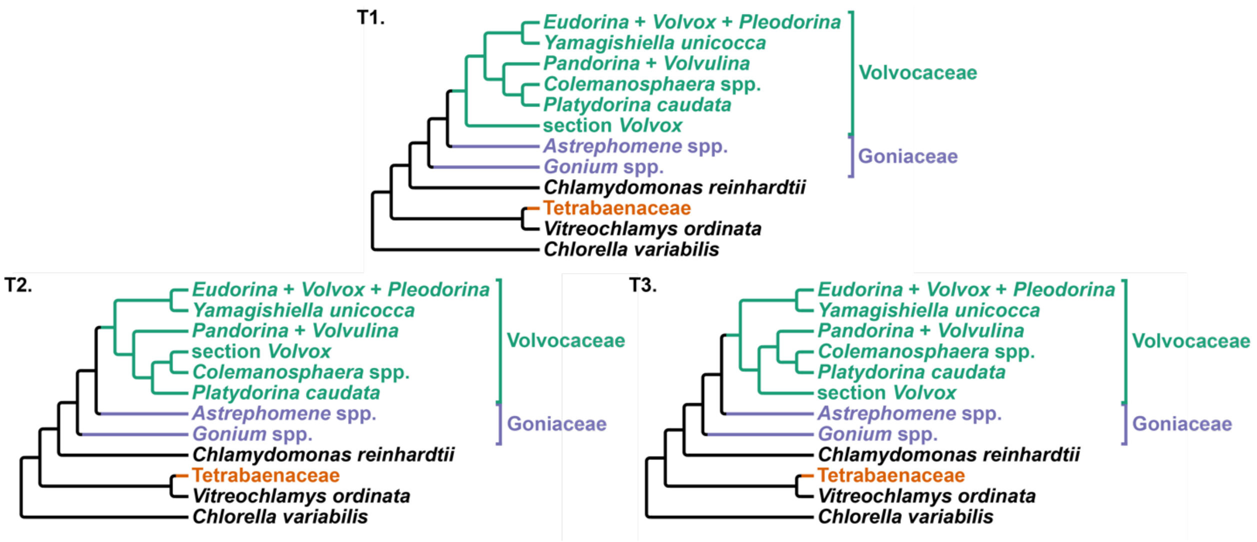


| Tree | logL | deltaL | p-AU |
|---|---|---|---|
| 1 | -308397.5847 | 0 | 0.967 + |
| 2 | -309045.8233 | 648.24 | 4.64e-88 - |
| 3 | -308434.3051 | 36.72 | 0.0332 - |

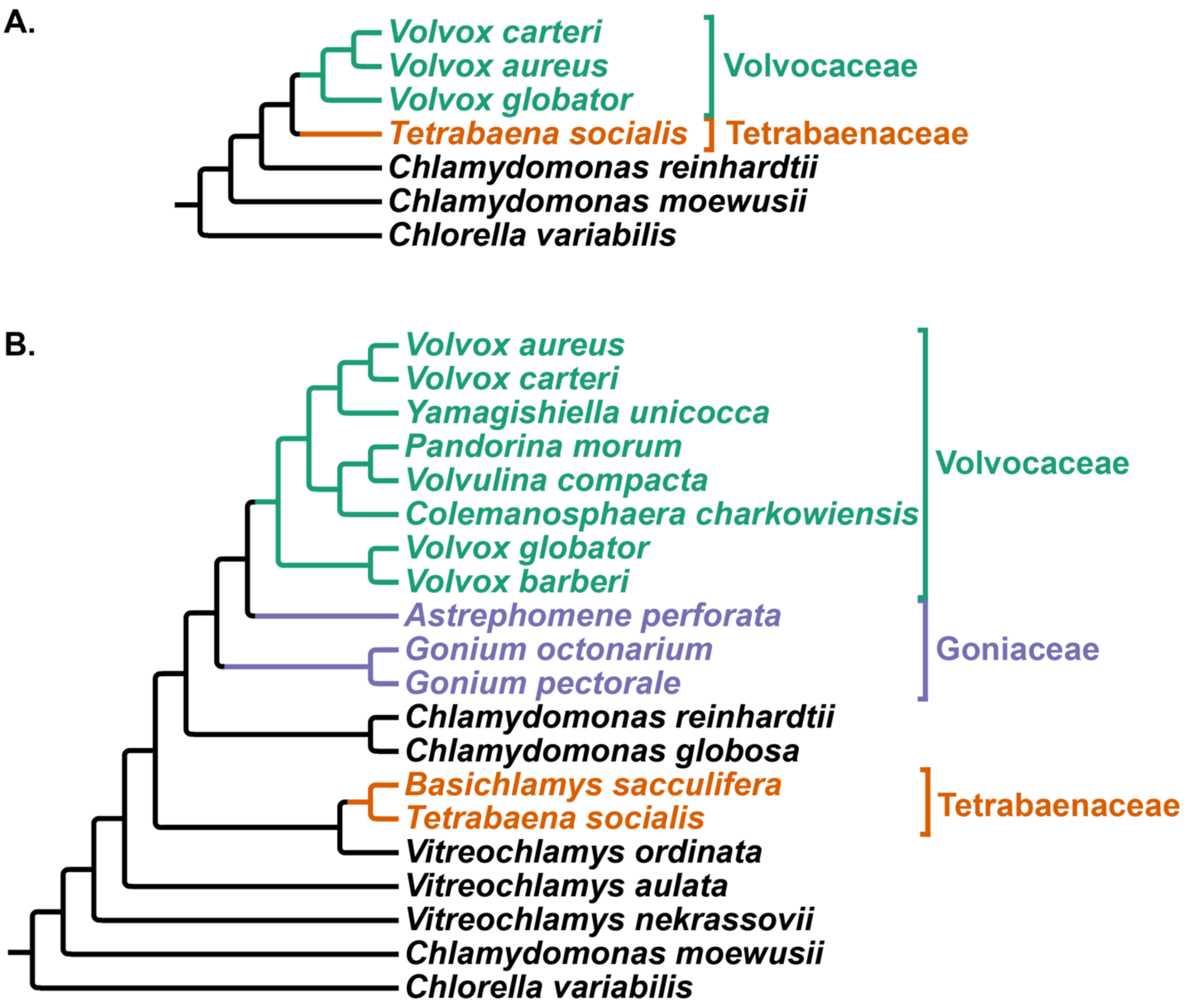


**Supplementary Figure S2. (A)** Phylogeny that represents a replication of the Zhang et al. (2019) results as they relate to the volvocine algae. The phylogenetic tree shown is based on 12,650 aligned amino acid positions among 7 taxa, and it was inferred using the maximum likelihood method in IQtree. **(B)** Phylogeny of the volvocine algae that represents a change in the branching order once more volvocine taxa are sampled for phylogenetic inference. This phylogeny shows the non-monophyly of the colonial algae and the Tetrabaenaceae sister to *Vitreochlamys ordinata*. The phylogenetic tree shown is based on 12,650 aligned amino acid positions among 20 taxa, and it was inferred using the maximum likelihood method in IQtree.

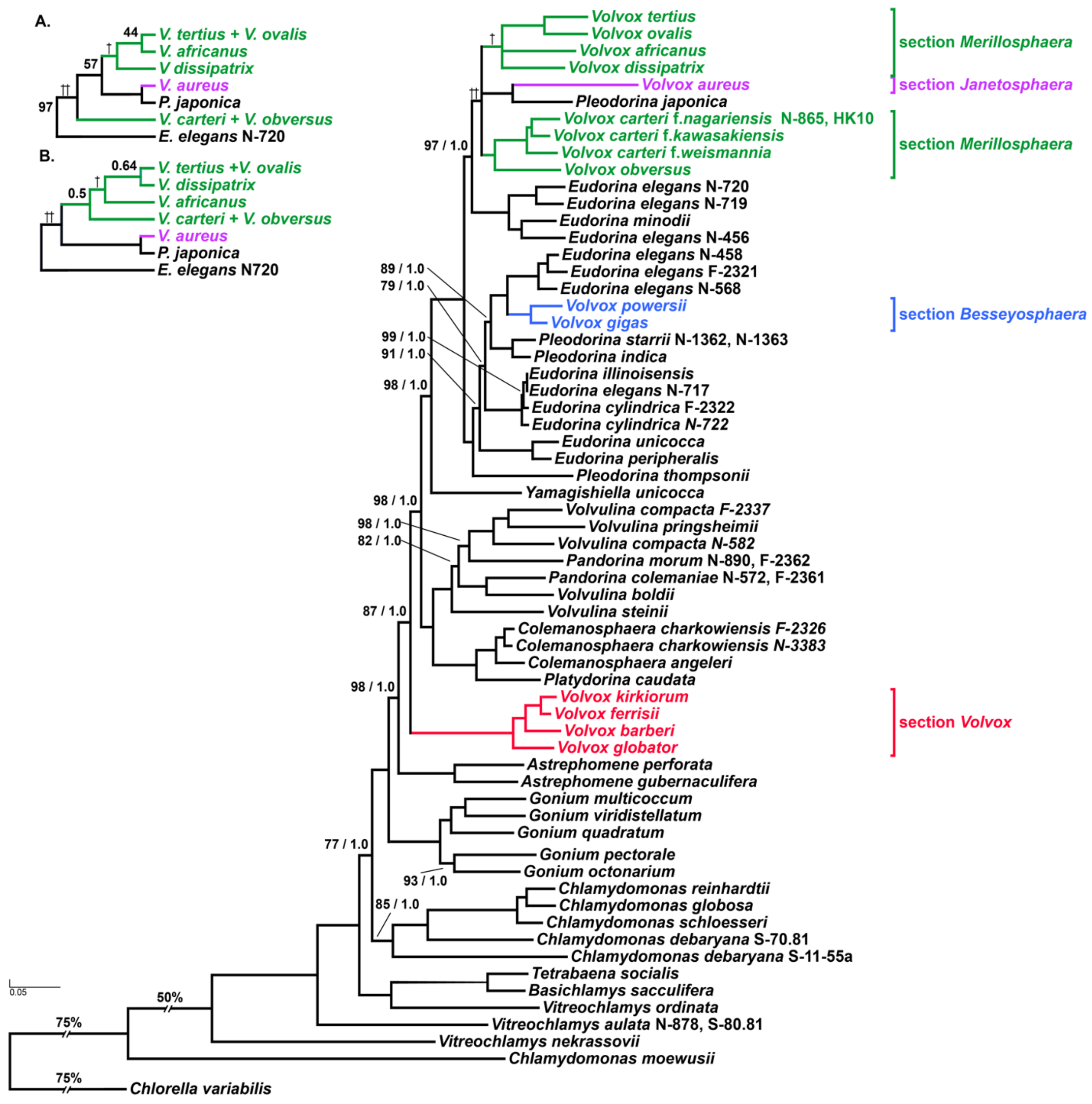


**Supplementary Figure S3.** Phylogeny of the volvocine algae which highlights the four sections of genus *Volvox* as recognized by Nozaki et al. (2015): *Besseyosphaera* (blue), *Janetosphaera* (purple), *Merrillosphaera* (green), and *Volvox* (red). The phylogenetic tree shown is based onv12,650 aligned amino acid positions of 68 taxa inferred using the maximum likelihood method in IQtree. Numbers on branches represent bootstrap values and Bayesian posterior probabilities, respectively. Branch lengths correspond to genetic divergence, as indicated by the scale bar. The Tetrabaenaceae, Goniaceae, and Volvocaceae are indicated by the orange, purple, and green highlight, respectively. Daggers correspond to topological differences between the maximum likelihood and Bayesian inference analyses that are represented in greater detail in Figures 2A and 2B, respectively.
