## Supplemental RNA Extraction Protocol Methods for "Phylotranscriptomics points to multiple independent origins of multicellularity and cellular differentiation in the volvocine algae"

***RNA extraction procedures*** Two RNA isolation protocols were followed for total RNA extraction: TRizol RNeasy and QIAGEN RNeasy Plant Mini Kit.

*TRIZOL RNEASY PROTOCOL* Cells/Colonies were pelleted and then resuspended in 3 mL 2% SDS in tris-buffered saline and 3 mL TRizol (guanidinium thiocyanate and acid phenol) in a 15 mL conical centrifuge tube and vortexed for approximately 30 seconds. Cells/Colonies were then flash frozen in liquid $N_2$. After sample was thawed, a phase separation was performed by centrifugation at 3000 *x g* for 10 minutes at room temperature. The aqueous phase was transferred to a 15 mL QIAGEN MaXtract High Density conical tube (prior to transferring the aqueous phase, the gel in the MaXtract tube was pelleted by centrifugation at 1500 *x g* for 1 minute). 0.6 mL chloroform was added to the MaXtract tube which was subsequently capped and shaken for approximately 15 seconds. After shaking, the MaXtract tube was left in a fume hood at room temperature for 3 minutes and then centrifuged at 1500 x g for 5 minutes at 4$^{o}$C to perform another phase separation. The aqueous phase was transferred to a 15 mL conical tube and 1.5 mL of isopropanol was added to the aqueous phase. The sample was left to incubate for 10 minutes at room temperature, and then another centrifugation step at 3000 *x g* for 10 minutes was performed. After centrifugation the supernatant was discarded, and the pellet was washed with 1 mL 75% ethanol. The ethanol was carefully pipetted off and the pellet was allowed to airdry for 5-10 minutes. The RNeasy mini spin column was centrifuged at 10,000 rpm or 10,976 x g for 15 seconds, followed by DNase digestion. Flow-through from the last step was discarded, and 350 μL of Buffer RW1 was added to the RNeasy Mini spin column and centrifuged at 10,000 rpm or 10,976 x g for 15 seconds. Flow-through was discarded. To a 70 μL aliquot of Buffer RDD was added 10 μL of QIAGEN RNase-free DNase. The resulting 80 μL Buffer RDD-DNase mixture was pipetted directly to the membrane of the RNeasy mini spin column and allowed to incubate for 15 minutes before proceeding with the protocol. After the incubation period, 350 μL of Buffer RW1 was added to the RNeasy spin column and centrifuged at 10,000 rpm or 10,976 x g for 15 seconds. Flow-through was discarded. 500 μL Buffer RPE was added to the RNeasy spin column and centrifuged at 10,000 rpm or 10,976 x g for 15 seconds. Flow-through was discarded. A second wash with 500 μL Buffer RPE was performed, and the RNeasy spin column was centrifuged at 10,000 rpm or 10,976 x g for 2 minutes. Flow-through was discarded, and the RNeasy spin column was placed inside a 1.5 mL collection tube supplied by QIAGEN. 30 μL of sterile RNase-free water was pipetted directly on the membrane and allowed to incubate for 1 minute, after which the RNeasy spin column was centrifuged at 10,000 rpm or 10,976 x g for 1 minute. RNA concentration and $A_{260}/A_{280}$ ratios were measured via Nanodrop (Thermo Fisher Scientific, Waltham, MA 02451 USA), and if RNA concentrations were >50 ng/μl then another elution step with 30 μL of RNase-free water was performed.

*QIAGEN PLANT MINI KIT PROTOCOL* Cells/Colonies were pelleted by centrifugation, resuspended in 500 μL of RLT Buffer RLT (β-mercaptoethanol was added to the Buffer RLT solution prior to starting) then vortexed for ca. 30 seconds. Resuspended cells/colonies were flash frozen in liquid $N_2$, then thawed and centrifuged at 10,000 rpm or 10,976 x g for 2 minutes in a QIAshredder spin column. The resulting lysate and any precipitate were transferred to a collection tube provided in the QIAGEN kit, whereafter 200 μL of 100% ethanol was added to the lysate and immediately mixed and transferred to an RNeasy Mini spin column. The RNeasy mini spin column was centrifuged at 10,000 rpm or 10,976 x g for 15 seconds, after which the optional DNase digestion step was performed. The flow-through from the last step was discarded, and 350 μL of Buffer RW1 was added to the RNeasy Mini spin column and centrifuged at 10,000 rpm or 10,976 x g for 15 seconds. Flow-through was discarded. A 70 μL aliquot of Buffer RDD was prepared and 10 μL of QIAGEN RNase-free DNase was added to it. The 80 μL Buffer RDD-DNase mixture was pipetted directly to the membrane of the RNeasy mini spin column and allowed to incubate for 15 minutes before the protocol was continued. Following incubation. 350 μL of Buffer RW1 was added to the RNeasy spin column, which was centrifuged at 10,000 rpm or 10,976 x g for 15 seconds. Flow-through was discarded. Then, 500 μL Buffer RPE was added to the RNeasy spin column and centrifuged at 10,000 rpm or 10,976 x g for 15 seconds. Flow-through was discarded. A second wash with 500 μL RPE buffer was performed, and the RNeasy spin column was centrifuged at 10,000 rpm or 10,976 x g for 2 minutes. Flow-through was discarded, and the RNeasy spin column was placed inside a 1.5 mL collection tube supplied by QIAGEN. 30 μL of RNase-free water was pipetted directly on the membrane and allowed to incubate for 1 minute. The RNeasy spin column was centrifuged at 10,000 rpm or 10,976 x g for 1 minute. RNA concentration and $A_{260}/A_{280}$ ratios were measured via Nanodrop. If RNA concentrations were >50 ng/μl, then another elution step with 30 μL of RNase-free water was performed.
