## Supplementary material for "Phylotranscriptomics points to multiple independent origins of multicellularity and cellular differentiation in the volvocine algae": translate_scaffold.txt

#!usr/bin/env python3

# The following script is used for a single taxon.
# Purpose of script: Read BLAST XML output files to get specific hit information (e.g. information regarding first BLAST hit, second BLAST hit, etc.), use that BLAST information to locate the scaffold of a gene we are wanting to find, translate the scaffold in all reading frames, and then add the entire scaffold in the appropriate reading frame to an alignment file. 
# Steps of the program:
# Step 1: Parse BLAST XML output files to get specific hit information. This program obtains three pieces of information from BLAST output: 1) "Query ID" which corresponds to the name of the query sequence 2) "Hit ID" which corresponds to the specific scaffold name in the transcriptome file where gene is located 3) "Hsp hit hseq" which corresponds to the sequence that best matches the query sequence.
# Step 2: Use the first hit id from BLAST XML file to locate the scaffold in the transcriptome file.
# Step 3: Translate the ENTIRE scaffold sequence in all three forward and all three reverse reading frames.
# Step 4: Locate the "Hsp hit hseq" in one of the 6 reading frames.
# Step 5: Write the original query name + query seq to a file with the hit organism name + hit sequence in a uniquely named directory.
# print commands (commented out) have been given throughout so that the process of the script can be seen in terminal.

import os
import xml.etree.ElementTree as ET  # Imports xml.etree.ElementTree which allows you to easily parse an XML file.
#  Regular expressions cannot (correctly) parse an XML file though I have seen it done. See famous post on StackOverflow for more information: https://stackoverflow.com/questions/1732348/regex-match-open-tags-except-xhtml-self-contained-tags/1732454#1732454
#  XML docs have sections called 'elements' which are defined by a beginning '<' and ending '>' tag.
#  The information contained within the tags '<info>' are called 'child elements'.
#  'root' is the main header at the top. It is called 'root' because the rest of the tree is rooted to it.
#  Subelements of the tree are called 'children'.
#  Sometimes tags will only contain text and you have to call it in your script as such.
#  If you want to navigate to a specific section of the XML file just type out the path.
from Bio.Seq import Seq  # Be sure to have biopython. Seq object allows you use Python strings with biological methods.
from Bio.Alphabet import generic_dna

path_to_vca_de_clerck = ""  # Path to Volvox carteri amino acid seqs that were shared with us by the De Clerck team.
vca_seqs_de_clerck = os.listdir(path_to_vca_de_clerck)

path_to_allxmltblastn_results = ""  # Path to the tblastn XML files that contain all BLAST hits for taxon.
xml_dir = os.listdir(path_to_allxmltblastn_results)

path_to_transcriptome = "" # PATH to appropriate transcriptome where scaffold ID will be located.
transcriptomefile = open(path_to_transcriptome, 'r')

output_dir = ""  # This creates the main output directory where the vca seqs and scaffolds translated in the correct reading frame will be output to.
if not os.path.exists(output_dir):  # This is checking to see if the path already exists. If it does not...
    os.makedirs(output_dir)  # ... then the directory that I named and assigned to a variable will be created.

# Read in tblastn results & identify top hit in tblastn results for all xml files (40) per species.
Query_Seq_id_list = []  # This list will be used later to create a dictionary: key= query_seq_id, value= first_hit_id
First_Hit_id_list = []  # This list will be used during the step where we search the transcriptome.
First_Hsp_hseq_list =[]  # This list will be used later when locating the correct reading frame translation.
for file in xml_dir:
    #print(file)
    if file.endswith(".xml"):
        xml_file_path = os.path.join(path_to_allxmltblastn_results, file)
        #print(xml_file_path)
        xml_files = open(xml_file_path, 'r')
        for lines in xml_files:
            #print(lines)
            tree = ET.parse(xml_files)  # ElementTree is now parsing all of the XML tblastn files.
            root = tree.getroot()  # This defines the root for each of the XML files. Since all the files have an identical format the root of each file is 'BlastOutput' (for my data, at least).
            #print(root.tag)

            for query_seq_id in root.findall(""):  # This for loop is capturing the query seq id that is associated to the De Clerck seq. In the XML file this corresponds to "<Iteration_query-def>" Under "<Iteration>".
                #print("query_seq_id =", query_seq_id.text)
                Query_Seq_id = query_seq_id.text
                Query_Seq_id_list.append(Query_Seq_id)

            for first_hit_id in root.findall(""):  # This for loop is capturing the Hit_id in the XML path of the first "Hit". This is specified by the "Hit[1] in the path. Example: "./BlastOutput_iterations/Iteration/Iteration_hits/Hit[1]/Hit_id"
                fill = first_hit_id.text  # Due to the format of the XML file the information contained within the tags are all 'text'. You must specify for the parser that the format of the info is 'text'.
                #fill2 = fill.replace('dbj', '')  # This and the replace below were added due to the Hit_id of one of the names having these characters added when they were not present in the normal tblastn results.
                #fill3 = fill2.replace('|', '')
                #First_Hit_id = fill3
                #print('First_Hit_id =', First_Hit_id)
                First_Hit_id_list.append(First_Hit_id)  # The for loop above where 'First_Hit_id' is captured only stores the final value when the program finishes its last loop. This step was necessary because we needed to store those values in a list in order to iterate over them.

            for first_hsp_hseq in root.findall(""):  # This for loop is capturing the first Hsp_seq in the path of the first Hit and first Hsp (this needs to be explicit because there can be multiple Hsp_hseqs in a BLAST hit). Example: "./BlastOutput_iterations/Iteration/Iteration_hits/Hit[1]/Hit_hsps/Hsp[1]/Hsp_hseq".
                filler = first_hsp_hseq.text
                replace_dashes = filler.replace('-', '')
                First_Hsp_hseq = replace_dashes
                #print('First_Hsp_hseq=', First_Hsp_hseq)
                First_Hsp_hseq_list.append(First_Hsp_hseq)

#print('first_hseq_list =', First_Hsp_hseq_list)

# Grab top hit scaffold ID from tblastn results and trace it back to the scaffold in the transcriptome.
frameF1_trans_list = []
frameF2_trans_list = []
frameF3_trans_list = []
frameR1_trans_list = []
frameR2_trans_list = []
frameR3_trans_list = []

for scaffold in transcriptomefile:
    if any(item in scaffold for item in First_Hit_id_list):
        contig_id = scaffold
        contig_nucl_seq = next(transcriptomefile)
        #print(contig_id)
        #print(contig_nucl_seq)

        # Translate the top hit scaffold in all 6 reading frames.
        # 5'-3' ORFs.
        scaffold_seq = Seq(contig_nucl_seq, generic_dna)
        #print("scaffold_seq =", scaffold_seq)
        frameF1 = scaffold_seq
        frameF2 = scaffold_seq[1:]
        frameF3 = scaffold_seq[2:]
        # print(frameF1)
        # print(frameF2)
        # print(frameF3)
        frameF1_trans = frameF1.rstrip().translate()
        frameF2_trans = frameF2.rstrip().translate()
        frameF3_trans = frameF3.rstrip().translate()

        frameF1_trans_list.append(frameF1_trans)
        frameF2_trans_list.append(frameF2_trans)
        frameF3_trans_list.append(frameF3_trans)

        #print("frameF1_trans =", frameF1_trans)
        #print("frameF2_trans =", frameF2_trans)
        #print("frameF3_trans =", frameF3_trans)
        # Reverse complement ORFs.
        complement_seq = scaffold_seq.complement()
        # print(complement_seq)
        rev_compliment = scaffold_seq.reverse_complement()
        # print("rev_compliment =", rev_compliment)

        frameR1 = rev_compliment[1:]
        frameR2 = rev_compliment[2:]
        frameR3 = rev_compliment[3:]

        # print("frameR1 =", frameR1)
        # print("frameR2 =", frameR2)
        # print("frameR3 =", frameR3)
        frameR1_trans = frameR1.rstrip().translate()
        frameR2_trans = frameR2.rstrip().translate()
        frameR3_trans = frameR3.rstrip().translate()

        frameR1_trans_list.append(frameR1_trans)
        frameR2_trans_list.append(frameR2_trans)
        frameR3_trans_list.append(frameR3_trans)

        #print("frameR1_trans =", frameR1_trans)
        #print("frameR2_trans =", frameR2_trans)
        #print("frameR3_trans =", frameR3_trans)
        # Use top tblastn hit to ID correct translation out of the 6 reading frames.

        six_frame_translation_list = [frameF1_trans_list, frameF2_trans_list, frameF3_trans_list, frameR1_trans_list, frameR2_trans_list, frameR3_trans_list]

correct_frame_translation_list = []
for frame in six_frame_translation_list:
    for item in frame:
        if any(sbs in item for sbs in First_Hsp_hseq_list):  # I used an arbitrary 'sbs' for 'substring' during this 'if statement' because I was finding a substring within a string (i.e. a partial sequence within a full sequence).
            #print('item =', item)
            correct_frame_translation_list.append(item)

#print('correct_frame_translation_list =', correct_frame_translation_list)
dictionary_Query_seq_and_Hit_id = dict(zip(Query_Seq_id_list, First_Hit_id_list))
dictionary_Hit_id_and_Hsp_hseq = dict(zip(First_Hit_id_list, First_Hsp_hseq_list))  # Dictionary contains keys (First_Hit_id) and its associated values (First_Hsp_hseq).
#print('dictionary =', dictionary_Hit_id_and_Hsp_hseq)

# This section takes the dictionary above (dictionary_Hit_id_and_Hsp_hseq) and replaces the incomplete First_Hsp_hseqs with the complete translated contig sequence.
for key, value in dictionary_Hit_id_and_Hsp_hseq.items():
    #print('key =', key)
    #print('value =', value)
    for trans_seq in correct_frame_translation_list:
        if value in trans_seq:
            dictionary_Hit_id_and_Hsp_hseq[key] = trans_seq
            #print('new_dic =', dictionary_Hit_id_and_Hsp_hseq)
            dictionary_Hit_id_and_correct_frame_translation = dictionary_Hit_id_and_Hsp_hseq

#print('dict_w_query_and_hit_ids =', dictionary_Query_seq_and_Hit_id)
#print('dict_w_correct_frame_translations =', dictionary_Hit_id_and_correct_frame_translation)

# Read into all the Volvox carteri seqs from De Clerck and append them to outfiles.
for file in vca_seqs_de_clerck:
    #print(file)
    if file.endswith(".fasta"):
        vca_seqs_path = os.path.join(path_to_vca_de_clerck, file)
        vca_seqs = open(vca_seqs_path, 'r')
        with open(os.path.join(output_dir, file), 'a') as output:
            for lines in vca_seqs:
                output.write(lines)

# Next two main For Loops are appending the correct scaffold name and its translated sequence in the correct reading frame to the same file with the De Clerck seq.
for filenames in os.listdir(output_dir):
    path_output = os.path.join(output_dir, filenames)
    #print(path_output)
    files_in_ouput = open(path_output, 'r')
    with open(path_output, 'a') as output2:
        for info in files_in_ouput:
            #print(info)
            for k, v in dictionary_Query_seq_and_Hit_id.items():
                if k in info:
                    output2.write(''+v+'\n')

for filenames in os.listdir(output_dir):
    path_output = os.path.join(output_dir, filenames)
    #print(path_output)
    files_in_ouput = open(path_output, 'r')
    with open(path_output, 'a') as output2:
        for info in files_in_ouput:
            for scaffold_id, translated_seq in dictionary_Hit_id_and_correct_frame_translation.items():
                if scaffold_id in info:
                    #print(str(translated_seq))
                    output2.write(str(translated_seq))

# Final step is to rename all files in output dir so that each retains their gene identifier but also states which organism each file belongs to.
for file_names in os.listdir(output_dir):
    new_filenames = file_names.replace('', '') # All files have '_aln' in name. This is finding it and replacing it with the species identifier and what the file is.
    os.rename(os.path.join(output_dir, file_names), os.path.join(output_dir, new_filenames))
